# Cytohesin-2 is essential for the survival of mice and regulates Golgi volume and function

**DOI:** 10.1101/2025.05.05.651226

**Authors:** Carsten Küsters, Bettina Jux, Farhad Shakeri, Sebastian Kallabis, Felix Meissner, Waldemar Kolanus

**Affiliations:** Department of Molecular Immune and Cell Biology, Life and Medical Sciences (LIMES) Institute, University of Bonn, Germany; Institute for Medical Biometry, Informatics and Epidemiology, Medical Faculty, University of Bonn, Bonn, Germany; Institute for Genomic Statistics and Bioinformatics, Medical Faculty, University of Bonn, Bonn, Germany; Systems Immunology and Proteomics, Institute of Innate Immunity, Medical Faculty, University of Bonn, Bonn, Germany; Core Facility Translational Proteomics, Institute of Innate Immunity, Medical Faculty, University of Bonn, Bonn, Germany

**Keywords:** Cytohesin-2, ARF-GTPases, Golgi apparatus, Protein secretion

## Abstract

Proteins of the cytohesin family are known for their guanine-nucleotide exchange factor function for ARF-GTPases, mainly for ARF1 and ARF6. While *Arf1* and *Arf6* deficiency results in embryonic lethality, *in vivo* functions of cytohesins are rarely described and mostly inconspicuous.

We analyzed the role of cytohesin-2 *in vivo* and *in vitro* and found that cytohesin-2 full knockout mice die within one day after birth. Mass spectrometry-based organellar proteomics in wildtype and CRISPR-Cas9-generated cytohesin-2^-/-^ C2 myoblasts revealed a markedly altered Golgi compartment. Golgi volumes were reduced in different cytohesin-2^-/-^ cell lines compared to wildtype cells as revealed by immunofluorescence. Reduced Golgi volumes were rescued by introducing cytohesin-2. Finally, we observed that typical functions of the Golgi apparatus were disrupted in cytohesin-2-deficient cells. Cytohesin2^-/-^ C2 myoblasts exhibited significant changes in the galactose / N-acetyl-galactosamine glycosylation on the cell surface compared to wildtype cells when stained with peanut agglutinin. Further, protein secretion was overall reduced in neonatal cytohesin-2^-/-^ mice compared to wildtype as determined by mass spectrometry-based proteomics.

This study describes the essential role of cytohesin-2 in neonatal development and a novel function of the protein in Golgi regulation.

## Introduction

ADP-ribosylation factor (ARF)–GTPases, depending on their cellular localization, are involved in processes such as membrane trafficking and organization of organelle structure (Gillingham and Munro 2007). Embryonic lethality of *Arf1-* and *Arf6*-deficient mice proves their crucial function in the organism (Hayakawa et al. 2014; Suzuki et al. 2006). The regulation of ARF activity, consisting of a cycle of GDP to GTP exchange by guanine-nucleotide exchange factors (GEFs) and GTP to GDP by GTPase activating proteins (GAPs), is therefore of importance as well. While large GEFs for ARF-GTPases such as GBP1 and BIG1/2 are mainly thought to be important for the organization of all parts of the Golgi apparatus, small GEFs are mainly located at the plasma membrane and endosomes. Proteins of the cytohesin (CYTH) family belong to the group of small GEFs and are therefore continuously discussed to be part of the vesicle transport machinery in the cell (Casanova 2007; Gillingham and Munro 2007). The CYTH protein family consists of four members of which CYTH1 and CYTH4 are mainly found in leukocytes while CYTH2 and CYTH3 appear to be ubiquitously expressed (Kolanus 2007). Their protein structure consists of an N-terminal coiled-coil domain that allows interaction between CYTHs and other proteins. Furthermore, CYTH proteins can be recruited to the plasma membrane by binding to the phosphatidylinositol-phosphates PtdIns(4,5)P_2_ and/or PtdIns(3,4,5)P_3_ via their C-terminal PH domain. Of note is the existence of two isoforms which differ in only one glycine residue within the PH domain. CYTH proteins in the two-glycine (2G) variant favor binding to PtdIns(3,4,5)P_3_, while the three-glycine (3G) variant mainly binds to PtdIns(4,5)P_2_. The GEF activity is mediated by the central sec7 domain (Chardin et al. 1996; Cronin et al. 2004; Klarlund et al. 2000). CYTH function can be inhibited by using the pan-CYTH inhibitor SecinH3 that binds to the sec7 domain and thereby blocks the GEF activity (Hafner et al. 2006). GEF-independent functions of CYTH proteins are barely described.

GEF activity of CYTH2 was mainly described for ARF1 and ARF6 and CYTH2 was found to regulate the cortical actin cytoskeleton in an ARF6-dependent manner, affecting epithelial cell migration (Santy and Casanova 2001; Santy et al. 2005). Additionally, CYTH2 was reported to regulate the endocytic route activating both ARF1 and ARF6, thereby facilitating Salmonella uptake, influenza virus transport and membrane trafficking towards the autophagosome (Humphreys et al. 2012; Hurtado-Lorenzo et al. 2006; Moreau et al. 2012; Yi et al. 2022). CYTH2 was also implicated in growth factor receptor signaling and its interaction with the Golgi apparatus was reported (Franco et al. 1998; Fuss et al. 2006; Hafner et al. 2006; Hurtado-Lorenzo et al. 2006; Jux et al. 2019; Lee and Pohajdak 2000; Monier et al. 1998). These findings derive from *in vitro* studies and the only *in vivo* reports using conditional knockout mice for *Cyth2* attributed the small GEF to neuronal activity and eosinic inflammation (Ito et al. 2021; Lolignier et al. 2015; London et al. 2022; Torii et al. 2015; Yamauchi et al. 2012).

To analyze the global role of *Cyth2* we have bred *Cyth2*-deficient mice, and we generated *Cyth2*-deficient cell lines by CRISPR-Cas9 to study CYTH2 function on vesicular transport in more detail *in vitro*. We found that all *Cyth2*-full knockout mice die within one day after birth. Overall reduced protein secretion due to a disturbed Golgi volume and function might explain this severe phenotype. With this study, we document a thus far unappreciated function for CYTH2 in regulating the Golgi apparatus.

## Results

### *Cyth2* full knockout leads to neonatal lethality

*In vivo* functions of CYTH proteins are to date sparsely investigated, therefore we bred full knockout mice of all four CYTH proteins. We and others observed no or minimal phenotypical effects in *Cyth1*, *Cyth3*, and *Cyth4* knockout mice without challenging (Jux et al. 2019; Yamauchi et al. 2012; and own unpublished data). In contrast, *Cyth2* full-knockout mice (heterozygous mice with conditional potential were obtained from EMMA) from heterozygous breeding were not detectable after weaning by genotyping although they were born in a Mendelian ratio (Fig. 1A). We observed that *Cyth2*^-/-^ mice died between 12 and 20 hours after birth (Fig. 1B). One to two hours after birth *Cyth2*^-/-^ mice had significantly less weight of 5.5% compared to wildtype (WT) littermates (Fig. 1C). Heterozygous littermates also had a slightly reduced body weight compared to WT mice and some pups died earlier (7 of 54) (Fig 1B, C) but *Cyth2*^+/-^ mice which grow to adulthood had no obvious phenotype. Besides the reduced weight we found that *Cyth2*^-/-^ mice never had a milk spot and were never observed to suckle (Fig. 1D) which could explain the weight loss. *Cyth2* is ubiquitously expressed (Suppl. Fig. 1A), and the highest expression level among tested organs was found in the brain. We generated several tissue-specific *Cyth2* knockout mice to identify the cause of death. Amongst others, we bred brain-specific (*Cyth2*^fl/fl^ x nestin-Cre) and skeletal muscle-specific *Cyth2* knockout (*Cyth2*^fl/fl^ x myogenin-Cre) mice. These tissue-specific knockout mice were vital and showed normal behavior without reduced life span (Suppl. Fig. 1B) and therefore excluded common neuromuscular defects as a reason for non-suckling and death. Kidney failure and respiratory defects are potential reasons for neonatal death. Mice with kidney failure can survive often more than 24 hours and Hematoxylin-Eosin-stained cryosections from six hours after birth looked inconspicuous (Suppl. Fig. 1C) (Turgeon and Meloche 2009). Also, lung sections from *Cyth2*^-/-^ mice looked comparable to WT controls in Hematoxylin-Eosin staining and severe lung defects usually lead to death within minutes (Suppl. Fig. 1D) (Turgeon and Meloche 2009). However, while we observed normal breathing activity in knockout mice during the first hours after birth, *Cyth2*^-/-^ pups developed cyanosis shortly before death. Thus, we cannot exclude respiratory defects as a cause of lethality. Furthermore, we generated heart-specific *Cyth2* knockout mice (*Cyth2*^fl/fl^ x alpha-Myosin Heavy Chain-Cre, *aMyHC*-Cre) which also did not phenocopy *Cyth2* full knockout mice (Suppl. Fig. 1B).

**Figure 1:**
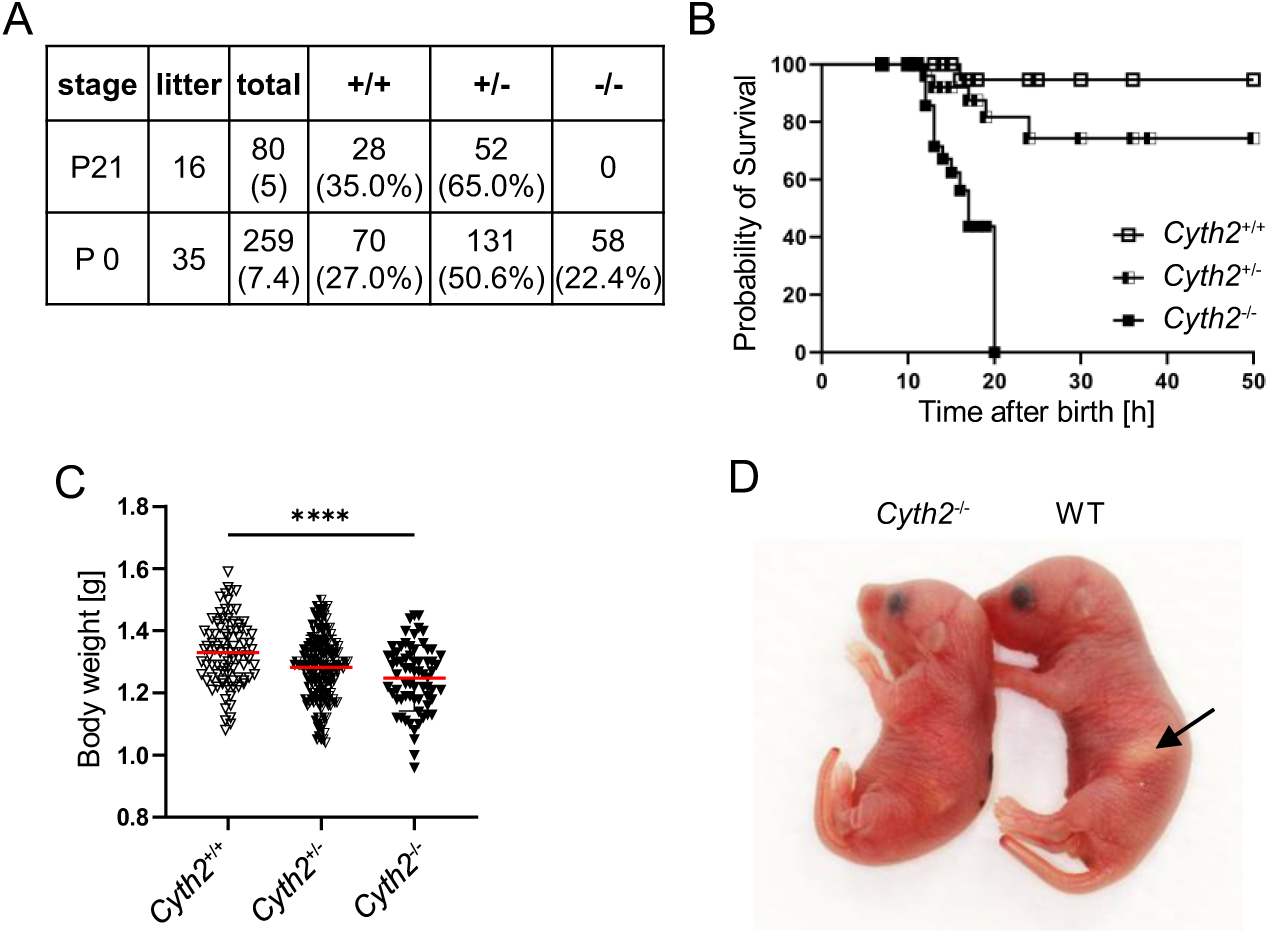
*Cyth2* full knockout leads to neonatal lethality. **A)** Percentage of viable *Cyth2* genotypes after weaning (P21) and newborn mice (P0). Given is the number of litters and mice which were analyzed. **B)** Survival rate of *Cyth2*^+/+^, *Cyth2*^+/-^ and *Cyth2*^-/-^ mice. Time of death was estimated as the midpoint between two timed observations. A total of 115 neonates were analyzed [*Cyth2*^+/+^ = 47 (1 died), *Cyth2*^+/-^ = 54 (7 died), *Cyth2*^-/-^ = 14]. **C)** Body-weight analysis of newborn pups one to two hours after birth. A total of 298 neonates were analyzed [*Cyth2*^+/+^ = 82, *Cyth2*^+/-^ = 151, *Cyth2*^-/-^ = 65, one-way ANOVA **** = P < 0.0001. **D)** Images showing *Cyth2*^+/+^ and *Cyth2^−^*^/−^ mice six hours after birth. In the Cyth2^+/+^ mouse, the arrow points to the milk spot, which is absent in *Cyth2^−^*^/−^ mouse.

Disturbed glucose homeostasis is another potential cause of death at time points we observed (Turgeon and Meloche 2009). Therefore, we generated and analyzed liver-specific (ls) knockout mice (*Cyth2*^fl/fl^ x albumin-Cre, ls *Cyth2*^-/-^). These mice were vital, with a normal life span, and showed no obvious abnormalities (Suppl. Fig. 2A). With food ad libitum WT and ls *Cyth2*^-/-^ mice had comparable weight and blood glucose level (BGL), normal expression of glucose homeostasis genes glucokinase (*Gck*), pyruvate kinase (*Pklr*), and phosphoenolpyruvate carboxykinase (*Pck1*), but a significant stronger expression of glucose-6-phosphatase (*G6pc1*) (Suppl. Fig. 2B-D). As insulin receptor signaling is decreased in *Cyth3*^-/-^ mice (Jux et al. 2019), we tested whether this is also the case in ls *Cyth2*^-/-^ mice. We found a significantly reduced phosphorylation of S6K after 10 minutes of insulin injection without differences in phosphorylation of AKT (Suppl. Fig. 2E-G).

An upstream kinase of S6K is mTOR, which has previously been described as an essential sensor for growth factors, amino acids, and other metabolites (Condon and Sabatini 2019; Saxton and Sabatini 2017). Hence, we analyzed BGL, serum amino acid level, and expression of glycolysis and gluconeogenesis genes in newborn *Cyth2*^-/-^ mice (Fig. 2). Although newborn *Cyth2*^-/-^ mice were not suckling they exhibited normal BGL compared to WT mice six hours after birth (Fig 2A). The BGL is tightly regulated by glycolysis and gluconeogenesis, we therefore tested for expression of *Gck* and *Pklr* in neonatal liver. However, these mRNAs were barely detectable in WT and *Cyth2*^-/-^ livers at six hours of age. The expression of gluconeogenesis genes *G6pc1* and *Pck1* was significantly upregulated in livers of *Cyth2*^-/-^ mice compared to WT which could explain the balanced BGL despite non-suckling (Fig. 2B). Serum amino acid levels are decreased in neonatal mice with impaired autophagy and mTOR signaling, respectively, and CYTH2-ARF6 activity was reported to promote autophagosome formation (Komatsu et al. 2005; Kuma et al. 2004; Moreau et al. 2012). Surprisingly, we found serum amino acid level increased in *Cyth2*^-/-^ mice compared to WT. Also, there was no indication for altered autophagy in *Cyth2*-deficient heart or lung tissue and observations in *Cyth2*^-/-^ myoblasts indicated even mildly increased autophagic activity (Fig, 2C; Suppl. Fig. 3). These findings hint at poor amino acid sensing and uptake in *Cyth2*-deficient animals.

**Figure 2:**
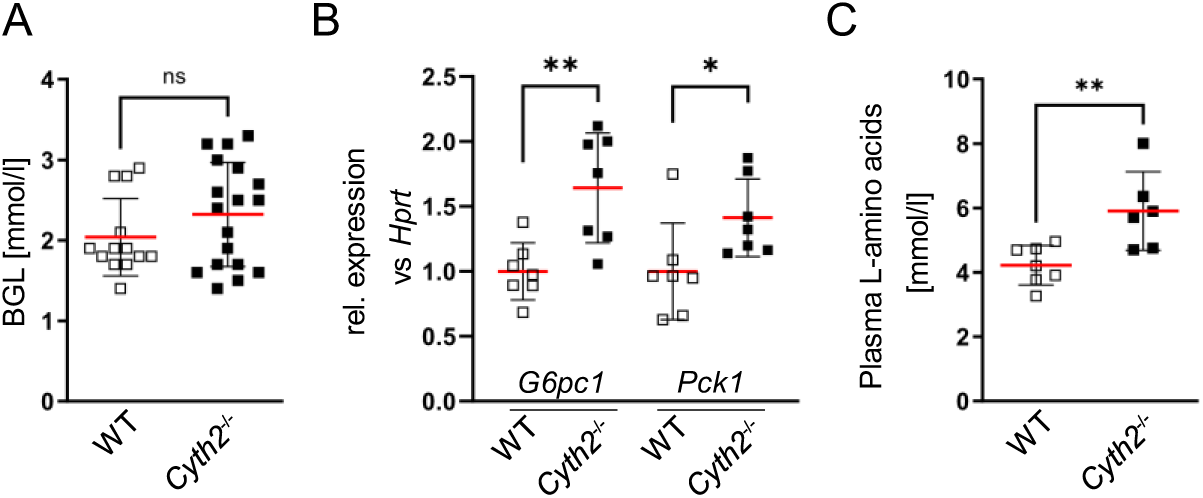
*Cyth2* full knockout mice have increased gluconeogenesis and plasma amino acids. **A)** Blood samples were taken from the neck after sacrificing the mice six hours after birth, n = 13 (WT) and 18 (*Cyth2*^-/-^). **B)** Gene expression of *G6pc1* and *Pck1* in livers from six hours-old WT and *Cyth2^−^*^/−^ (KO) mice was analyzed by PCR. The expression was normalized to *Hprt*. **C)** L-amino acids were measured in plasma isolated from six hours old mice. Results are given in means ±SD. Every dot represents the result of a single mouse. (*p < 0.05; **p < 0.01; ns = not significant).

Taken together, *Cyth2* is essential for neonatal survival and involved in metabolic regulations.

### *Cyth2*^-/-^ myoblasts have decreased ULK1 phosphorylation upon stimulation with amino acids

To analyze the role of *Cyth2* in the sensing of metabolites in more detail we generated *Cyth2*^-/-^ C2 myoblasts by CRISPR-Cas9. We used C2 myoblasts as they are fast-growing, easy to handle, metabolically active, and no tumor cells. After proving efficient knockout by western blot (Fig. 3A) we performed re-feeding experiments by first depriving WT and *Cyth2*^-/-^ myoblasts of amino acids for two hours, followed by restimulation with full medium containing amino acids. We analyzed downstream signaling, namely phosphorylation of AKT and the mTOR targets S6K and ULK1 (Fig. 3A). As expected, phosphorylation levels of S6K and ULK1 were low in starved cells and strongly induced after the addition of amino acids. However, while phosphorylation of S6K reached comparable levels between WT and Cyth2^-/-^ cells, phosphorylation of UK1 was significantly lower in knockout myoblasts. AKT phosphorylation on Thr308 was not affected by amino acid starvation and re-feeding and was independent of *Cyth2*, indicating unhindered growth factor receptor signaling (Fig. 3A). We also tested the impact of glucose and FCS starvation and restimulation on C2 myoblasts (Suppl. Fig. 4). Deprivation and restimulation of C2 cells with glucose had no effect on the phosphorylation of ULK1, S6K, and AKT and phosphorylation was independent of *Cyth2* (Suppl. Fig. 4A). FCS starvation abolished S6K and AKT phosphorylation completely and strongly reduced ULK1 phosphorylation (Suppl. Fig. 4B). Addition of FCS induced phosphorylation of all three target sites tested in WT and *Cyth2*^-/-^ cells with a slight but not significant reduction of ULK1 phosphorylation in *Cyth2*^-/-^ cells. These results indicate a selective defect in mTOR target phosphorylation for ULK1 in *Cyth2*-deficient C2 myoblasts in response to re-supplementation of amino acids.

**Figure 3:**
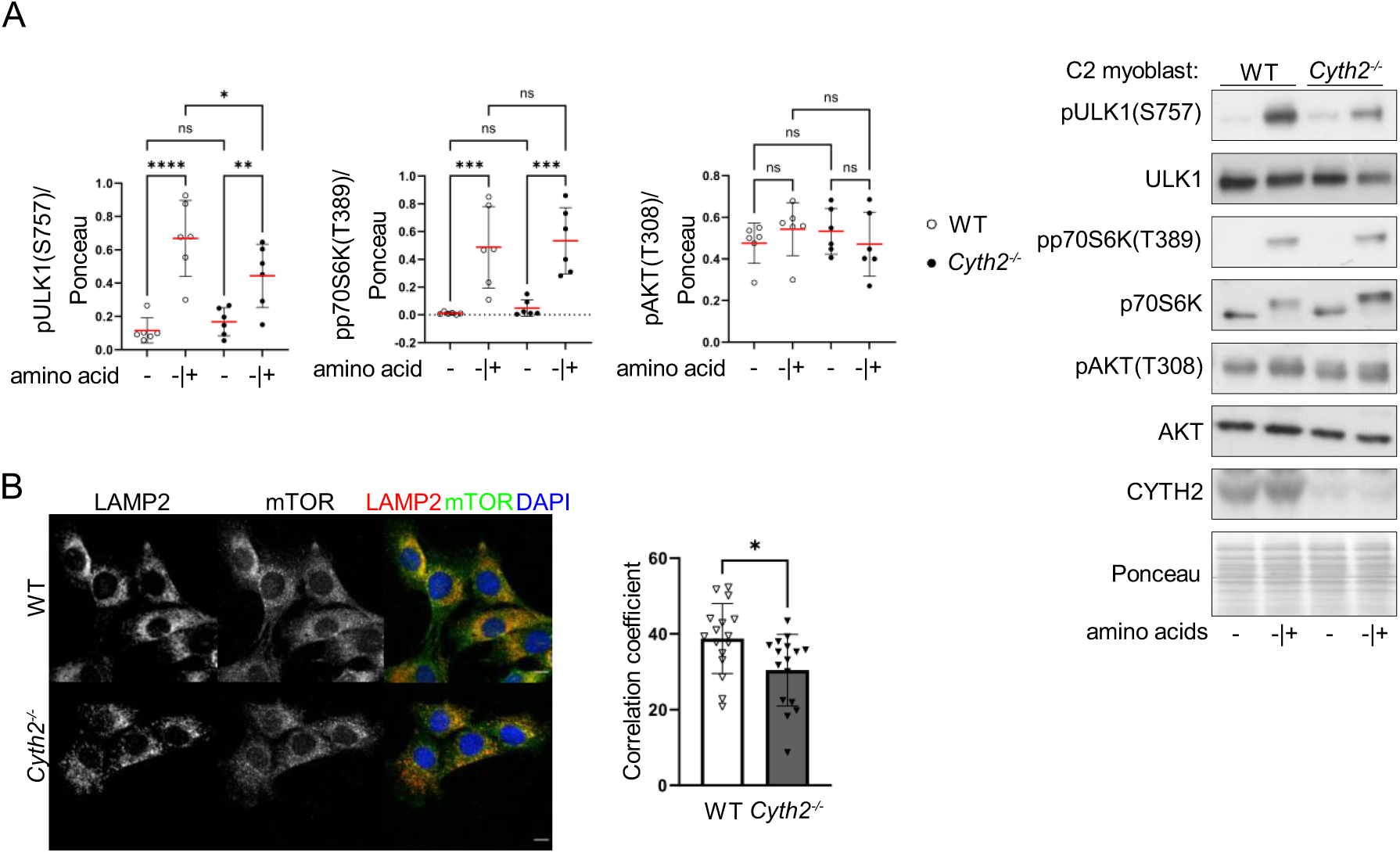
Reduced phosphorylation of the mTOR target ULK1 and reduced recruitment of mTOR to lysosomes in *Cyth2*^-/-^ myoblasts. **A)** Western Blot analysis of phosphorylated ULK1, p70S6K and AKT after deprivation of amino acids for two hours (-) and subsequent restimulation (-|+) for two hours. Left: Average signal intensity (±SD; n=6), normalized to total protein loaded as determined by Ponceau staining. Right: representative immunoblot analyses are shown. **B)** Lysosomal localization of mTOR analyzed by co-staining with LAMP2 after two days standard culture. Bars indicate mean Pearson’s correlation coefficient (±SD) for mTOR and AMP2 calculated with Imaris 9. Data were pooled from four experiments, including two independent pairs of WT and *Cyth2*^-/-^ clones. Bars represent mean±SD, each dot represents one z-stack analyzed (n=16, 16). Significance was calculated using 2way-ANOVA and unpaired t-test (*p < 0.05; **p < 0.01; ***p < 0.001; ****p < 0.0001; ns = not significant).

Activation of mTOR involves the relocation of the mTORC1 complex to the surface of lysosomes and defective recruitment prevents effective downstream signaling (Sancak et al. 2010). Therefore, we analyzed the mTOR recruitment to lysosomes by co-staining of the kinase with the lysosomal marker LAMP2 (Chen et al. 1985) in WT and *Cyth2*^-/-^ C2 myoblasts under normal culture conditions (Fig. 3B). Interestingly, we found a small but significant reduction of mTOR-LAMP2 co-localization in *Cyth2* knockout cells.

Taken together, we find reduced phosphorylation of the mTOR target ULK1 and reduced recruitment of mTOR to lysosomes in *Cyth2*^-/-^ myoblasts compared to WT C2 cells which again hints at reduced amino acid sensing in these cells.

### Organellar proteome maps reveals changes in Golgi structure

As CYTH2 has been implicated in vesicle transport processes (Casanova 2007), we asked whether a loss of *Cyth2* might hinder the correct positioning of amino acid transporters, thereby affecting the activation of mTOR and its downstream signaling towards UlLK1. We chose an unbiased approach to answer this question and performed differential centrifugation of WT and *Cyth2*^-/-^ C2 myoblasts to separate organelles by their sedimentation properties. The resulting fractions were subjected to spectrometry-based proteomics analysis and the abundancies of proteins in each fraction were compared between WT and KO to identify displaced proteins and altered organelles (Itzhak et al. 2019). Principle component analysis (PCA) separates different cellular compartments by specific marker proteins, previously described as organellar maps (Itzhak et al. 2016) (Fig. 4A). Overall, the PCA plots were comparable between WT and *Cyth2*^-/-^ cells, showing that overall cellular integrity is not disturbed by the knockout of *Cyth2*. We detected 177 proteins, which were significantly displaced in *Cyth2*^-/-^ C2 myoblasts compared to WT cells in two independent experiments (Fig. 4B). These proteins clustered in two groups and Fisher’s exact test was performed to identify significantly enriched GO Terms in these two clusters, among which we identified “signal-anchor” as the strongest enriched term (Fig. 4C). Comprising many glycosylating enzymes, this finding prompted us to analyze organellar markers, and indeed a disturbed distribution of Golgi markers in *Cyth2*^-/-^ cells compared to WT cells was visible with a clear displacement from the 5500xg fraction towards fractions of lower and higher centrifugation speed (Fig. 4D, F). We also identified displacement of marker proteins of other cellular compartments, such as the plasma membrane, actin-binding, and endosomes, but to a lesser extent.

**Figure 4:**
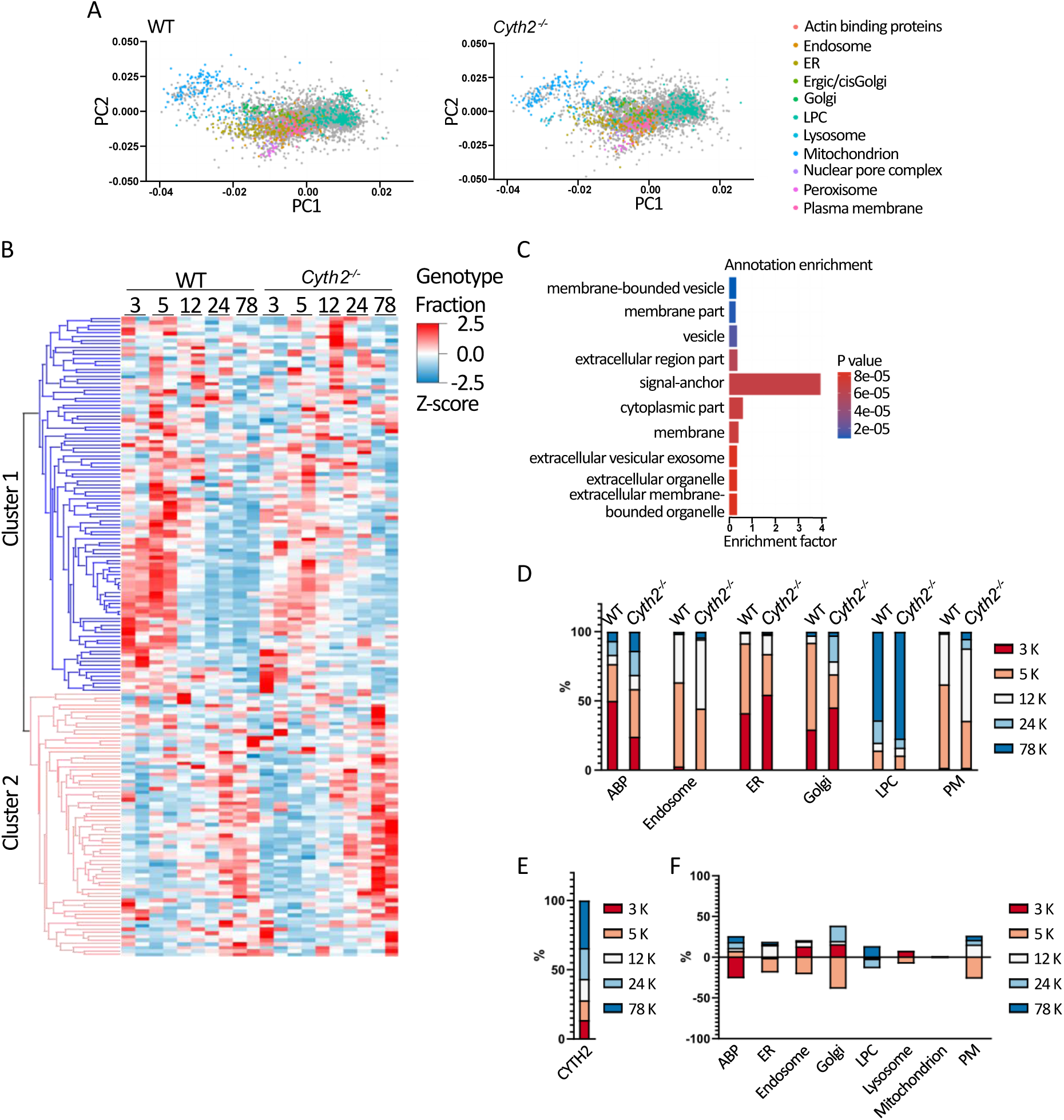
Dynamic organellar proteome maps reveal a displacement in Golgi-associated proteins. **A)** Representative PCAs of proteins detected in differential centrifugation fractions of WT (left) and *Cyth2*-deficient (right) myoblasts based on their LFQ intensities. Marker proteins as identified by Itzhak (Itzhak, 2016) are depicted in color according to the legend, grey dots represent all other proteins. **B)** Heatmap of significantly displaced proteins in both experiments, assigned to two clusters (unsupervised). **C)** Fisher’s exact test for GO term enrichment in the two clusters in B). **D)** Percentage of major organellar proteins (as in A) across the five fractions, exemplarily shown for actin-binding proteins (ABC), endosomes, the ER, Golgi, large protein complexes (LPC), and the plasma membrane (PM). **E)** WT-KO difference of percentages of marker proteins in E).

The localization of CYTH2 within a cell has not been fully elucidated so far due to a lack of specific antibodies. Different studies find the majority in the cytosol (Frank et al. 1998; Li et al. 2007), the plasma membrane (Frank et al. 1998; Torii et al. 2010; Venkateswarlu et al. 1998), endosomes (Maranda et al. 2001; Salem et al. 2015), and the Golgi compartment (Lee and Pohajdak 2000). The localization might be dependent on the status of the cell. And the functions at different sites are not well understood but accepted for the endosomal pathway. Employing organellar maps, we were able to confirm, that CYTH2 is localized at endosomes and the plasma membrane, but we also found CYTH2 enriched in protein complexes (Fig. 4E).

Our organellar maps of *Cyth2*^-/-^ C2 myoblasts compared to WT controls support a function for CYTH2 in vesicular transport of the endocytic pathway, because endosomal markers were displaced in *Cyth2*-deficient C2 myoblasts. Additionally, a novel role of CYTH2 in regulating Golgi-associated vesicle transport appears possible.

### Loss of *Cyth2* reduces Golgi volume in different cells from different species

To validate the findings from the protein localization analysis, we exploited immunofluorescence analysis. We confirmed changes in the endosomal compartment, as staining C2 myoblasts and A7r5 cells for RAB5 revealed increased numbers of early endosomes in *Cyth2*^-/-^ cells compared to WT controls (Suppl. Fig. 5). The accumulation of early endosomes was propagated to the late endosomal compartment as revealed by increased numbers of RAB7-positive structures in knockout cells (Suppl. Fig. 5). In accordance, we observed elevated numbers of lysosomes (LAMP2-positive) earlier in *Cyth2*^-/-^ C2 myoblasts compared to WT cells (Suppl. Fig. 3F). While previous studies addressing a role of Cyth2 in endocytosis focused on the localization of cargo proteins such as the transferrin receptor, we analyzed RAB5 and RAB7 as key regulators of endocytosis (Naslavsky, 2023).

However, perturbation of the Golgi compartment as a consequence of *Cyth2* ablation was unexpected, and represents a novel aspect of *Cyth2* biology, since the integrity of the Golgi compartment was so far mainly attributed to the function of large ARF-GEFs such as GBP1 and BIG1/2 (Casanova 2007; Gillingham and Munro 2007). CYTH2 was shown before to have some minor localization to the Golgi apparatus in Cos-1 cells (Lee and Pohajdak 2000), and it acts as a GEF for ARF1 (Chardin, 1996; Cohen, 2007; Hafner, 2006; Humphreys, 2012), but a regulation of the Golgi compartment by CYTH2 has not been explored. To validate morphological perturbation of the Golgi apparatus by ablation of CYTH2 function, we stained the Golgi complex in WT and *Cyth2*^-/-^ C2 myoblasts with the common Golgi marker Golgin-97 (also Golga1), and evaluated the organelle structure using IMARIS software (representative pictures Fig. 5A). We found that overall Golgi volume was reduced in two independent CRISPR-Cas9 *Cyth2*^-/-^ clones of C2 myoblasts compared to WT control cells (Fig 5B). To validate this further, we generated *Cyth2*^-/-^ HEK 293T cells (human, kidney, Fig. 5C) and A7r5 cells (rat, smooth muscle, Fig. 5D) with CRISPR-Cas9. The *Cyth2* knockout was verified by western blot and Golgi structure was assessed by immunofluorescence staining of Golgin-97. Comparable to C2 myoblasts the deletion of *Cyth2* in HEK 293T cells (Fig. 5C) and A7r5 cells (Fig. 5D) also led to significantly reduced Golgi volumes.

**Figure 5:**
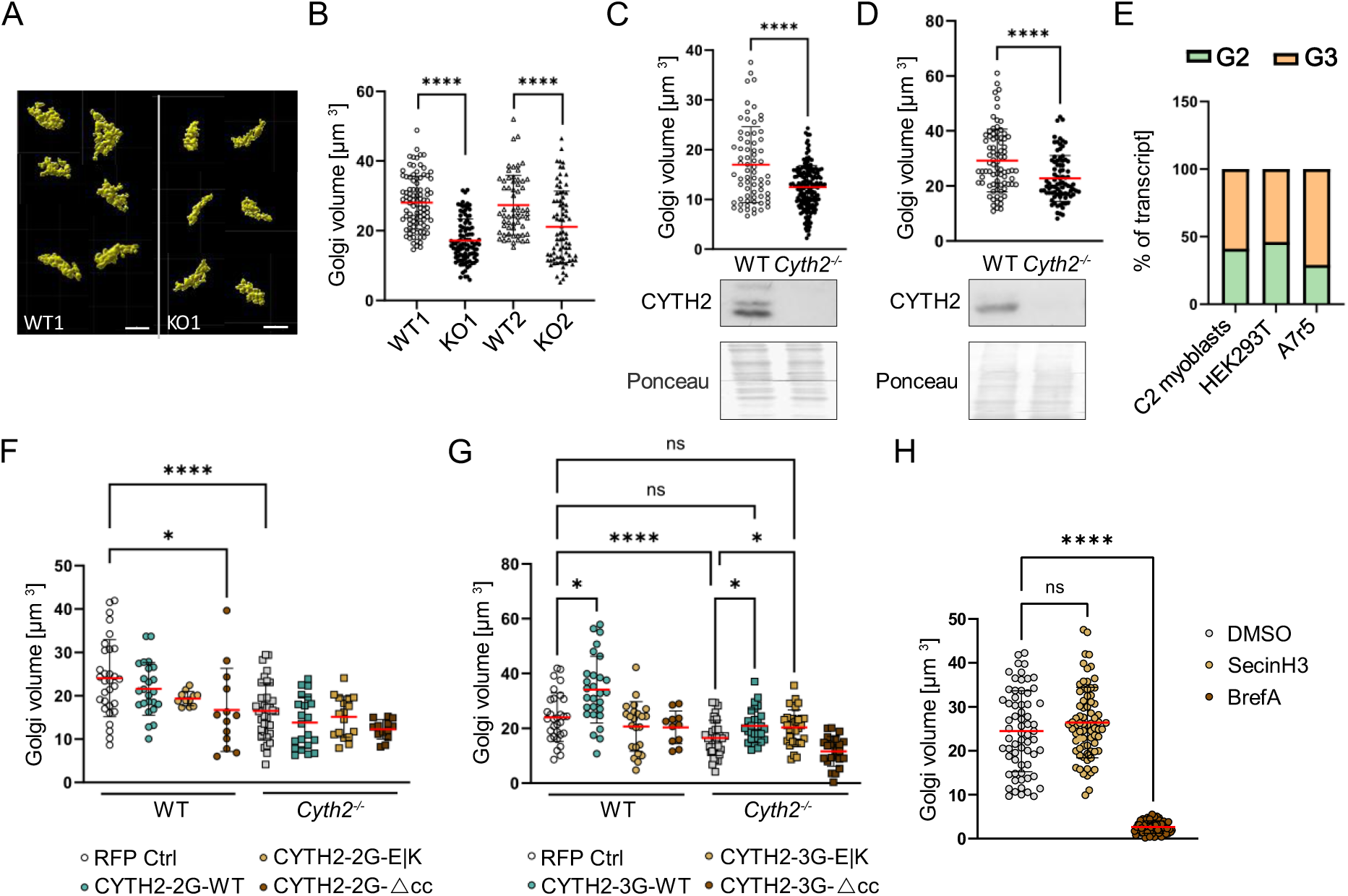
Loss of *Cyth2* reduces Golgi volume in different cells from different species and can be rescued by CYTH2 overexpression. **A)** Examples of reconstructed Golgis based on z-stack confocal images of Golgin-97-stained WT and *Cyth2*-deficient myoblasts detected with Imaris 9 surface detection. **B-D)** Golgi volume analyzed by Imaris 9 in B) two pairs of WT and *Cyth2*-knockout C2 myoblasts clones, as well as in WT and *Cyth2*-deficient C) HEK 293T and D) A7r5 cells under standard culture conditions. In addition, C-D) include Western Blot controls for the depletion of *Cyth2*). **E)** Quantified expression of the two *Cyth2* isoforms by Sanger sequencing and Tide analysis in WT C2 myoblasts, HEK 293T and A7r5 cells. **F-G)** Golgi volume analyzed as in B-D) in WT and *Cyth2* knockout myoblasts overexpressing CYT 2 WT, E|K and Δcc in the F) 2G and G) 3G isoform, RFP serving as a control. **H)** Golgi volume analyzed as in B-D) in WT C2 cells treated with 50 µM SecinH3 or 5 µg/ml Brefeldin A for 4 h, DMSO serving as control. Each symbol represents one Golgi apparatus analyzed; red lines indicate means±SD (pooled data from more than three independent experiments. Significance was tested using Mann-Whitney (B-D)), 2way-ANOVA (F-G)) or One-way ANOVA (H)). In F-G) only relevant and significant comparisons are depicted (*p < 0.05; **p <0.01; ***p < 0.001; ****p < 0.0001; ns = not significant).

Taken together, CYTH2 is involved in the structural regulation of the Golgi apparatus in different cells and organisms.

### Golgi volume reduction can be rescued by CYTH2-3G in an ARF-GEF-independent fashion

CYTH2 is expressed in two isoforms, differing in a single glycine residue within the PH domain. To define, which isoform is responsible for the observed regulation of Golgi volume, we first determined the *Cyth2* variant expressed in C2 myoblasts, A7r5 cells and HEK293T cells. Sanger Sequencing of amplified *Cyth2* transcripts isolated from WT cells revealed expression of both, the 2G (∼40%) and 3G isoform (∼60%) (Fig. 5E). This ratio was comparable between the different cell types analyzed. Therefore, rescue experiments were performed in C2 myoblasts using both isoforms. We furthermore included the GEF-inactive mutant (E156K; E|K) and a CYTH2 construct lacking the coiled-coil domain (Δcc), which should result in its reduced binding capacity to other proteins and recruitment to the Golgi apparatus (Lee and Pohajdak 2000; Mansour et al. 2002; Nevrivy et al. 2000; Venkateswarlu 2003). C2 myoblasts were transfected with fusion proteins of the mentioned CYTH2 variants (2G, 3G, E|K, Δcc) with RFP or an RFP-expressing vector alone as a control. Then, Golgin-97 was stained and the structure of the Golgi apparatus was analyzed by fluorescence microscopy in RFP-positive cells. RFP-transfected *Cyth2*^-/-^ myoblasts showed decreased Golgi volumes compared to RFP-expressing WT controls (Fig. 5F, G), confirming previous results and showing, that the transfection method has no effect on Golgi volume. Expression of the 2G variant of CYTH2 could not reverse the decreased Golgi volume in *Cyth2*^-/-^ C2 myoblasts, neither the wildtype nor the CYTH2-2G-E156K mutant or the CYTH2-2G-Δcc variant (Fig. 5F). In contrast, introduction of wildtype CYTH2-3G in *Cyth2*^-/-^ C2 cells was able to rescue the reduced Golgi volume (Fig. 5G). Also, in WT C2 cells CYTH2-3G increased the Golgi volume over the levels of RFP-transfected WT cells. GEF-inactive CYTH2-3G (E|K) likewise rescued Golgi volumes in knockout myoblasts, while the CYTH2-3G-Δcc variant was unable to do so. Treatment of WT C2 cells with SecinH3, a small molecule inhibitor of the CYTH GEF function did not change Golgi volume in contrast to Brefeldin A, which caused disassembly of the Golgi complex (Fig. 5H). This finding supports that the effect of CYTH2 on the Golgi volume is independent of its GEF function, at least in this *in vitro* system.

Taken together, these data show a clear involvement of CYTH2-3G in Golgi volume regulation, while the 2G variant has no effect on Golgi volume. Furthermore, Golgi volume controlled by CYTH2-3G appears to be independent of ARF activation but might depend on protein binding of CYTH2.

### *Cyth2* deficiency reduces Golgi functions such as glycosylation and secretion

The Golgi apparatus plays a major role in posttranslational modification of transmembrane and secreted proteins, involving the attachment of sugar moieties to the protein known as glycosylation (Joshi et al. 2018; Marth and Grewal 2008; Schjoldager et al. 2020). These sugar residues form a broad range of antigens on proteins with different complexity, which are recognized by lectins, making lectins a biochemical tool to quantify these glycosylated residues. We therefore analyzed the glycosylation status of WT and *Cyth2*^-/-^ C2 myoblasts, labelled cells with fluorophore-labeled lectin, and subjected them to flow cytometric analysis (Fig 6A). We stained with mannose-binding lectin (Concanavalin A, ConA), galactose / N-acetyl-galactosamine-binding lectin (Peanut agglutinin, PNA), and N-acetyl-glucosamine-binding lectin (Wheat germ agglutinin, WGA). In all cases, cell surface tracing of protein glycosylation was significantly reduced by treatment of wildtype C2 cells with the Golgi inhibitor Brefeldin A, showing that these glycosylation patterns are regulated by the Golgi apparatus (Fig. 6A, B). While binding of ConA and WGA was unaffected by *Cyth2* ablation, PNA labeling was significantly reduced by 40% in two independent CRISPR-Cas9 generated *Cyth2*^-/-^ C2 cell clones (Fig. 6A, B), clearly showing a defect during the glycosylation process within the Golgi. This result was supported by the finding, that the C1GALT1-C1GALT1C1 complex mediating galactose / N-acetyl-galactosamine modifications is displaced in the absence of CYTH2 as determined in the organellar maps analysis (Fig. 6C). Further, in the organellar maps we found the amino acid transporter SLC7A5 significantly displaces (Fig. 6D). This finding not only hints to a probable defect in amino acid sensing in *Cyth2*^-/-^ mice (leading to high plasma L-amino acid level) and C2 cells (leading to a reduced mTOR localization and downstream signaling), it also shows that Golgi-mediated protein transport to the plasma membrane might be disturbed as well.

**Figure 6:**
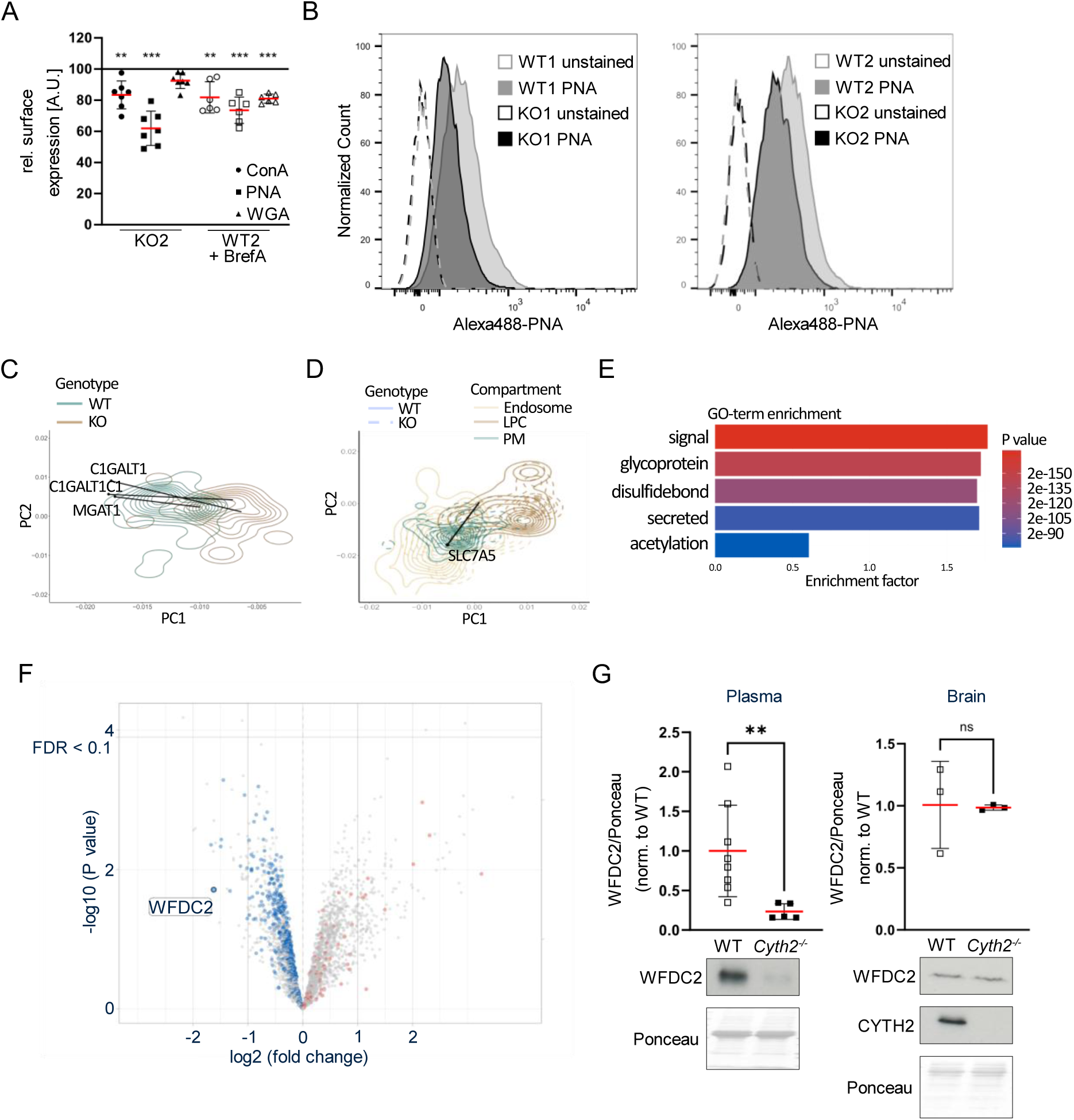
*Cyth2*^-/-^ C2 cells have a reduced galactose / N-acetyl-galactosamine glycosylation, altered positioning of the amino acid transporters SLC7A5 and *Cyth2*^-/-^ mice have an impaired secretion. **A)** Relative MFI of lectin staining (ConA, PNA, WGA) for one clone pair of C2 myoblasts comparing *Cyth2*-deficient or Brefeldin A-treated WT cells to untreated WT controls (100% reference line), measured by FACS. Each dot represents one independent experiment (n ≥ 6), red lines represent mean±SD. **B)** Representative histograms of PNA measured in two pairs of C2 myoblast clones as in A). **C)** PCA-based density plots of the Golgi compartment (extracted from Fig. 4A) overlayed with PC1 and PC2 of C1GALT1, C1GALT1C1 and MGAT1 in WT (black dot) and *Cyth2* knockout (end of black line) myoblasts. **D)** PCA-based density plots of endosomes, large protein complexes (LPC) and plasma membrane (PM) overlayed with PC1 and PC2 of SLC7A5 as in C), for WT and KO. Block dot represents the SlLC7A5 WT PCs, the end of the black line the KO PCs. **E)** GO-term enrichment analysis (Fisher’s exact for down-regulated proteins) of proteins measured in plasma from neonatal WT and *Cyth2*-deficient mice by proteomics. **F)** Volcano plot (of proteomics data analyzed in F), comparing KO/WT. Secreted proteins (GO: KW-0964) are highlighted in blue (down) and red (up). All other proteins are represented as grey dots. **G)** Western Blot analysis of WFDC2 in neonatal plasma (left) or brain (right) from WT and *Cyth2*-deficient mice. Top: Average signal intensity (red line±SD; n≥3) normali ed to total protein loaded as determined by Ponceau staining. Bottom representative immunoblot analyses are shown. Significance was tested with the Mann-Whitney test (*p < 0.05; **p < 0.01; ***p < 0.001; ns = not significant).

As we here and others show that a reduced Golgi function leads to a false or reduced glycosylation (Ahat et al. 2022; Ignashkova et al. 2017; Merk et al. 2009; Misumi et al. 1986). This in turn could lead to a failure in the protein secretion process. Therefore, we asked, whether we find fewer secreted proteins in the plasma of neonatal *Cyth2*^-/-^ mice compared to wildtype mice. We analyzed plasma samples from six hours old mice by proteomics. The GO-term enrichment analysis of down-regulated proteins revealed terms of glycosylation and secretion as significantly enriched (Fig. 6E). The volcano plot highlights that secreted proteins (colored in blue and red) are overall reduced in the plasma of *Cyth2*^-/-^ mice (Fig. 6F).

One of the severely reduced proteins in neonatal plasma of knockout animals was WFDC2 (WAP Four-Disulfide Core Domain Protein 2). WFDC2 is a secreted glycoprotein and it was recently shown that *Wfdc2*-deficiency leads to neonatal lethality (Drapkin et al. 2005; Kirchhoff et al. 1991) (Nakajima et al. 2019; Zhang et al. 2020). We therefore examined WFDC2 in plasma from six hours old mice by western blot and confirmed the proteomics findings: In plasma from *Cyth2*^-/-^ mice WFDC2 was significantly reduced compared to plasma from WT and heterozygous mice. In contrast, in neonatal brains the expression level was comparable between WT and knockout mice, highlighting an effect of *Cyth2* on WFDC2 secretion, not its expression (Fig. 6G). Of note, due to its glycosylation secreted WFDC2 in the plasma has a higher molecular weight of appr. 27 kDa compared to WFDC2 that resides in the tissue with a molecular weight of appr. 18 kDa (Drapkin et al. 2005). As *Cyth2*^-/-^ mice do not entirely phenocopy *Wfdc2*^-/-^ we assume that neonatal death of *Cyth2*^-/-^ mice is attributed to diminished secretion of several proteins.

Taken together, these data indicate that CYTH2 regulates Golgi volume and function in various cell types. The absence of CYTH2 leads to impaired glycosylation and a failure in protein secretion.

## Discussion

In our organellar maps analysis, we found a displacement of many proteins, among them ∼20% of endosomal marker proteins. We confirmed changes in the endo-lysosomal compartment, finding the numbers of early and late endosomes as well as lysosomes increased in cells lacking CYTH2 (Suppl. Fig. 3F, 5). Our results shed new light on aspects of CYTH2 regulating the endocytic process, since previous studies had analyzed CYTH2-dependent endocytosis focusing on cargo protein transport alone, but data on the functional interaction of CYTH2 with regulatory proteins such as the RAB GTPases are lacking. The range of studies addressing varying aspects of endocytosis and a regulatory role of CYTH2 in the process emphasizes the need for a comprehensive study of CYTH2-dependent endocytosis, observing cargo, regulatory proteins and effector proteins at the same time.

The unbiased analysis of subcellular protein localizations in *Cyth2*-deficient cells has revealed a novel aspect of *Cyth2* biology and identified many Golgi-associated proteins to be displaced (Fig. 4). Furthermore, we could subsequently confirm changes in Golgi structure via immunofluorescence analysis, and found the Golgi volume to be significantly reduced in *Cyth2*^-/-^ C2 myoblasts, A7r5 and HEK293T cells compared to WT controls (Fig. 5). Moreover, the Golgi volume could specifically be rescued by introducing CYTH2 into knockout cells. CYTH2 has GEF activity for ARF1, the major ARF-GTPase regulating Golgi function (Gillingham and Munro 2007; Pennauer et al. 2022; Sztul et al. 2019) but the effect of CYTH2 on Golgi volume in C2 cells appears to be GEF-independent (Fig. 5G, E156K). Possibly, CYTH2 plays a role as an adaptor protein while for instance the large GEFs activate ARF1 (Casanova 2007; Gillingham and Munro 2007). Additionally, based on cell-free transport assays, an ARF-independent transport between Golgi cisternae was proposed (Happe and Weidman 1998). Therefore, it seems more likely that CYTH2 affects the Golgi apparatus influencing the localization of proteins possibly due to interaction with the coiled-coil domain. This is underlined by the finding, that the effect on Golgi volume could not be rescued by a cc-deficient CYTH2 mutant. That CYTH2 was able to rescue the Golgi volume when specifically overexpressed in the three-glycine isoform points to the relevance of PtdIns(4,5)P_2_ in this context and emphasizes the validity of our findings. As described before “PtdIns(4,5)P_2_ may contribute to the structural organization of the Golgi by interacting with cytoskeletal elements” (Mayinger 2009).

Golgi architecture and integrity are essential for posttranscriptional modifications and transport of proteins throughout the cell, including protein secretion. Activation of ARF1 by ARF-GEFs plays an essential role in cargo sorting and vesicle formation (Adarska et al. 2021; Arakel and Schwappach 2018). While large GEFs such as GBF1 and BIG1/2 are essential to maintain Golgi integrity and function, small GEFs with a C-terminal PH domain such as CYTHs are described to act at the plasma membrane and endosomes (Casanova 2007; Gillingham and Munro 2007; Sztul et al. 2019). Even though a localization of CYTH2 to the Golgi apparatus depending on the coiled-coil domain was shown before (Lee and Pohajdak 2000), investigations about its function at the Golgi are missing.

The present study suggests a structural and functional impact of CYTH2 on the Golgi apparatus in several cell types. Among the proteins displaced in the organellar maps we found C1GALT1 which regulates glycosylation. In accordance with that we found a reduced PNA binding on the surface of *Cyth2*^-/-^ C2 myoblasts compared to WT cells. Furthermore, we found the amino acid transporter SLC7A5 displaced. It was shown before, that the knockdown of SLC7A5 led to a reduced mTOR activity (Sokolov et al. 2020). We see a reduced mTOR activity and lysosome localization in *Cyth2*^-/-^ C2 cells compared to WT C2 cells. This is in accordance to a recent publication, that describes *Cyths* as necessary for proper mTOR localization to lysosomes after amino acid stimulation (Luo et al. 2021). The delocalization of SLC7a5 could also explain the higher plasma amino acid level in *Cyth2*^-/-^ mice compared to WT mice, as the uptake of amino acids is disturbed (Nässl et al. 2011).

Failures in Golgi assembly are the cause of many diseases and knockout of Golgi proteins leads to diverse phenotypes, ranging from early lethality to infertility and others (Jiang et al. 2020; Kim et al. 2020; McGee et al. 2017; Zahn et al. 2006). We found fewer secreted proteins in the plasma from *Cyth2*^-/-^ mice compared to WT mice, which were not attributable to specific organs. One of the proteins was WFDCc2. Recently it was shown, that *Wfdc2*^-/-^ mice die 10 hours after birth due to a lung failure. CYTH2 is highly expressed in the lung and we cannot exclude lung defects as a cause of death. However, as *Cyth2*^-/-^ mice do not entirely phenocopy *Wfdc2*^-/-^ mice though, we assume that neonatal death of *Cyth2*^-/-^ mice is potentially attributed to overall reduced Golgi functionality, including diminished secretion of various proteins, reduced glycosylation and disturbed localization of Golgi-passing proteins such as the amino acid transporter SLC7A5.

So far, only three *in vivo* studies have analyzed conditional knockout mice of *Cyth2*. Brain-specific *Cyth2*^-/-^ mice (Ito et al. 2021) show reduced mechanical allodynia in inflammatory and neuropathic pain models but otherwise, these mice were normal in lifespan. This confirms our observation, that although CYTH2 is highest expressed in the brain, brain-specific knockout mice show no phenotype without challenge. Also, a Schwann cell-specific knockout of *Cyth2* causes no aberrant phenotype despite leading to a weak myelinization defect (Torii et al. 2015). The most recent publication describes a reduced eosinophilic inflammation in conditional *Cyth2* mice which were crossed with a germ cell-specific Cre line (London et al. 2022). However, it cannot be excluded, that the knockout via Cre-recombination was not fully efficient and controls are missing, which might explain a rather mild phenotype of that particular mouse line. The *Cyth2*^-/-^ mice of our study are indeed *Cyth2* full knockout mice as has been proven by western blot (Suppl. Fig. 1A). Apart from reduce body weight, *Cyth2* knockout mice show normal appearance, movement, breathing, and no malformations. By breeding tissue-specific knockout mice we exclude for instance neuromuscular or hepatic defects as a cause of death. Also, non-suckling seems unlikely to be a direct effect of the *Cyth2* knockout, but a secondary effect e.g., due to a general weakness deduced from the reduced weight of *Cyth2*^-/-^ mice directly after birth. Our observations with various tissue-specific knockout mice emphasize, that only a full knockout of *Cyth2* seems to have a severe effect and hints to a more systemic defect in full knockout animals.

Heart-specific *Cyth2*-deficiency is no exception, yet needs to be mentioned here: Cardiomyocyte-specific *Cyth2*^-/-^ mice died 8 – 12 months after birth due to a dilated cardiac hypertrophy (Suppl. Fig. 6A-F). However, it turned out to be an effect of the *aMyHC*-Cre line and seemed to be independent of the *Cyth2* expression (Suppl. Fig. 6G). The *aMyHC*-Cre line (Torii et al. 2015; Agah et al. 1997) is therefore not recommended to generate a stable heart-specific knockout (Li et al. 2023).

In the present study, we first describe that *Cyth2*-deficiency leads to a reduced Golgi volume in different cell lines with functional consequences for glycosylation, protein transport to the plasma membrane, and potentially protein secretion in mice. The latter provides a plausible explanation for the neonatal lethal phenotype of *Cyth2* full knockout mice, yet requiring further investigation. Together, we could show that *Cyth2* contributes to Golgi structural and functional integrity and is essential for the survival of mice.

## Material and Methods

### Mice

*Cyth2*^+/-^ mice (B6NDen;B6N-Cyth2^tm1a(EUCOMM)Wtsi/Ibcm^) were obtained from European Mouse Mutant Archives (EMMA, Rome, Italy). Male and female C57BL/6N wildtype (WT) and *Cyth2*^-/-^ mice originate from heterozygous breeding pairs and were bred under specific pathogen–free conditions at the animal facility of the University of Bonn. Mice were sacrificed by decapitation at the age of six hours for removal of organs and plasma. Animal care and experiments were performed according to German and Institutional guidelines for animal experimentation and were approved by the government of North Rhine-Westphalia (Germany).

### Genotyping

Genotyping of neonatal mice was done from the tail tip. For retraceability mice were marked on the back with a number, which had no impact on the maternal behavior. DNA was isolated by incubation of tissue material with 200 µl 50 mM NaOH for 20 min at 95 °C. The reaction was stopped by adding 70 µl Tris/HCl pH 8 and subsequent centrifugation at 4000 rpm for 4 min. Primer sequences are given in Suppl. Table 1.

### Blood sample handling

Blood samples were taken from neck after decapitation of neonatal mice or from the tail tip of adult mice and blood glucose was determined with the Accu-Chek^®^ system (Roche, Mannheim, Germany). 20 µl of blood was collected in a heparinized Microvette^®^ CB300 (Sarstedt, Nümbrecht, Germany). After centrifugation, the plasma was transferred into a fresh tube and stored at −80 °C. For determination of plasma L-amino acids the L-Amino Acid Quantification Kit from Merck (Darmstadt, Germany) was used according to the manufacturer’s instruction.

### Real time PCR

Total RNA was isolated with TRIzol^™^ and transcribed into cDNA using the High-Capacity cDNA Reverse Transcription Kit (Applied Biosystems™, CA, USA). Real-time PCR was performed on an CFX96^™^ Real-Time System (Biorad, München, Germany) using the iTaq™ Universal SYBR^®^ Green Supermix (Biorad). Expression levels were normalized to *Hprt* as housekeeping gene (IDT, Leuven, Belgium; primer sequences are listed in Suppl. Table 1).

### Cell culture

C2 myoblasts were cultured in DMEM medium supplemented with 15% FCS, 2% sodium pyruvate, 1% non-essential amino acids, 100 U/mL penicillin and 0.1 mg/mL streptomycin (P/S). For all nutrient restimulation experiments dialyzed FCS (10 kDa cut-off; Invitrogen/Life Technologies) was used. Cells were deprived for the respective nutrient for 2 h and restimulated with full medium for another 2 h. HEK 293T cells were cultured in DMEM medium containing 10% FCS and P/S. A7r5 cells were cultured in DMEM medium without phenol red and low glucose, supplemented with 10% FCS, P/S and 2% L-Glutamine (all cell culture components from PAN-Biotech, Aidenbach, Germany).

### CRISPR-Cas9 knockout of *Cyth2* and rescue

Knockout cell lines were generated using the CRISPR/Cas9 system combined with lentiviral transduction as described before (Joung et al. 2017). gRNAs were used as published in the Human and Mouse CRISPR Knockout Pooled Library (Brunello and Brie library; Suppl. Table 2 (Doench et al. 2016)) for gene editing. gRNA oligonucleotides were cloned into the pLentiCRISPRv2 vector (Transfer plasmid, Addgene, #52961) using BsmBI restriction and Golden Gate assembly. gRNA-containing pLentiCRISPRv2 plasmids were co-transfected with the pMD2.G envelop plasmid (Addgene, #12259) and psPAX2 packaging plasmid (Addgene, #12260) into HEK 293T cells using the calcium phosphate transfection method and incubated for 48 h to produce lentivirus particles. Filtered supernatant (0.45 µm pore size) was supplemented with 8 µg/ml polybrene and added to the target cell lines for 24 h. After additional 24 h transduced cells were selected with puromycin for three days, followed by subcloning (for C2 and HEK cells). Effective gene editing was confirmed by Western Blot and Sanger Sequencing for single-cell clones.

For rescue experiments, C2 myoblast clones were seeded on cover slips. After cells adhered, they were transfected with the appropriate plasmids (Suppl. Table 3) with jetOPTIMUS® (Polyplus transfection®). After 48 hours cells were fixed and stained for analysis of the Golgi apparatus.

### Western Blot analysis and antibodies

Cells and organs from WT and *Cyth2*^-/-^ mice were homogenized on ice in MRC-lysis buffer (50 mM Tris-HCl, pH 7.5, 1 mM EGTA, 1 mM EDTA, 270 mM Sucrose, 1% Triton X-100, protease- and phosphatase-inhibitors). Western blots were performed as described previously (Mitschka et al. 2015). After blotting, nitrocellulose membranes were stained with Ponceau Red to determine equal loading of samples. Antibodies: CYTH2 (clone H-7, Santa Cruz), GAPDH (ACP001P, Acris), pAKT (T308; #9275), AKT (#9272), p p70S6K (T389; #9205), p70S6K (#9202), pULK1 (S757; #6888), LC3b (#2775), (all Cell Signaling Technologies) and WFDC2 (PA5-80227).

### Lectin staining and flow cytometry

C2 myoblasts were harvested with 2 mM EDTA in PBS. Cells were stained with Alexa Flour 488 conjugated Lectins (Thermo Fisher: Con A-Alexa488 conjugate C11252; PNA-Alexa488 conjugate L21409, WGA-Alexa555 conjugate W32464) and measured with a FACS Canto II. Analysis was done with FlowJo™ (v.10).

### Differential centrifugation and mass spectrometry

The differential centrifugation to analyze subcellular protein localization was performed as described by Itzhak et al (Itzhak et al. 2016). Briefly, C2 cells were harvested and washed once in ice-cold hypotonic lysis buffer (HLB, 25mM Tris pH 7.5, 50mM Sucrose, 0.2 mM EGTA, 0.5 mM MgCl_2_ and protease inhibitors). Cells were lysed in HLB in a dounce homogenisator with 20 strokes. Cell homogenate was immediately mixed with 10% v/v hypertonic sucrose buffer (25 mM Tris pH 7.5, 2.5 M Sucrose, 0.2 mM EGTA, 0.5 mM MgCl_2_ and protease inhibitors) to restore the sucrose concentration to 250 mM. The cell lysate/supernatant was sequentially centrifuged at 1000g, 10 min.; 3000g, 10 min.; 5500g, 15 min.; 12200g, 20 min; 24000g, 20 min.; 78400g, 30 min. The pellet was resuspended in RIPA-buffer (150 mM NaCl, 1% TritonX, 0.5% NaDoc, 0.1% SDS, 50 mM Tris pH7.5). Samples were stored at −20 °C until analysis.

Proteins enriched in organellar fractions as well as from total cell lysate were denatured using urea lysis buffer (6 M urea, 2 M thiourea, 10 mM Tris-(2-carboxyethyl)-phosphine, 30 mM 2-chloroacetamide in 50 mM Tris-HCl pH 8.5). After incubation at room temperature for 30 min, urea concentrations were diluted to less than 1 M using 50 mM Tris-HCl. Proteins were enzymatically digested at room temperature with Trypsin/Lys-C (Promega) at a 1:100 enzyme-to-protein ratio for 16 hours. On the next day, the digestion was stopped by adding 10% formic acid to a final concentration of 1% and peptides were desalted using a modification of the stage tip protocol (Rappsilber et al. 2007). In-house-made polystyrene-divinylbenzene reversed phase sulfonate (SDB-RPS, Affinisep) double-layer stage tips were activated by the addition of 100 µl pure methanol. Tips were washed with 100 µl buffer B (80 % acetonitrile, 0.1% formic acid in LC-MS-grade water) and equilibrated with 100 µl buffer A (0.1% formic acid in LC-MS-grade water). Digested peptides were loaded onto the stage tips with a volume equivalent to 20 µg of initial protein input material and following were desalted by one wash of 100 µl buffer A and two washes of 100 µ buffer B. Elution of peptides was achieved by the addition of 60 µl of buffer X (1% ammonia in acetonitrile). After each step, centrifugation was carried out at 800 g for two minutes or until all liquid passed through the double layer. Finally, eluted peptides were vacuum-dried, and dissolved in buffer R (2% acetonitrile, 1% formic acid in LC-MS-grade water), peptide concentrations were determined, and 400 ng was used for injection into the LC-MS system.

Proteomics measurements were carried out with an ultrahigh-performing liquid chromatography Easy nLC 1200 system coupled on-line to an Orbitrap Exploris 480 tandem mass spectrometer (both Thermo Fisher). Peptides were chromatographically separated on an in-house produced analytical column (30 cm length, 75 µm inner diameter, 1.9 µm ReproSil-Pur 120 C18-AQ filling material [Dr Maisch]) and using a 90 min gradient consisting of buffer A and B. Starting at 4% the amount of Buffer B was linearly increased to 25% over 70 minutes at a flow of 300 nl/min. Buffer B was then linearly increased to 55% over 8 minutes and finally to 95% within 2 minutes. The analytical column was washed at 95% B for another 10 min.

Eluting peptides were on-line-transferred into the MS system by nanoelectrospray ionization operated at a constant voltage of 2.4 kV. Samples were analyzed in data-independent acquisition (DIA) mode. In brief, full MS spectra were recorded at a resolution of 120,000, a scan range of 380 – 1,020 m/z, an AGC target of 100%, and an injection time of 55 ms. DIA fragment spectra were recorded at a resolution of 15,000, an AGC target of 1,000% and an injection time of 22 ms. In total, 75 DIA windows of 8 m/z window sizes were selected spanning a range from 400 – 1,000 m/z. Precursors were fragmented at an HCD collision energy of 31% and measured in centroid mode.

Mass spectrometry raw data were processed with the DIA-NN software tool (Demichev et al. 2020). A spectral library was predicted *in silico* from the UniProt SWISS-PROT Mus musculus database (version from 2021-11-18) with trypsin as the digesting enzyme, maximum number of missed cleavages set to 1 and cysteine carbamidomethylation enabled as a fixed modification. The scan window radius was set to 7 and mass accuracies were fixed to 2.08e-05 (MS2) and 3.6e-06 (MS1), respectively. Precursor masses were fixed to min 380 m/z and max 1,020 m/z with peptide sequence lengths from 7 – 30. Peptide-spectra matches were filtered at an FDR < 0.01. Identified precursors were further filtered in R on Lib.Q.Values and Lib.PG.Q.Values < 0.01. Label-free quantified (LFQ) protein intensities were carried out with the maxLFQ algorithm implemented in the DIA-NN R package (Cox et al. 2014).

The statistical analysis was performed with the Perseus software suite (Tyanova et al. 2016) (v. 1.6.15). The following steps were performed consecutively to identify proteins which changed organellar localization upon protein knockout. Firstly, summed intensities were calculated for each protein within wildtype or knockout separately. Summed intensities were then subtracted individually from intensities in each fraction to get relative protein abundances. Delta abundances were calculated per fraction by subtracting KO relative abundances from WT relative abundances to reveal proteins changing organellar localization upon genotype. Only proteins present in each fraction of KO and WT were considered for multidimensional significance testing (Threshold = 0.05, Quantile = 0.55, iterations = 100, Benjamini-Hochberg FDR). Proteins with significantly different localizations were Z-score normalized and used for hierarchical clustering (Euclidean distance, clustering = k-means, 300 starting points, 10 iterations). Finally, enriched gene ontology terms enriched in individual clusters compared to all proteins were identified by Fisher’s Exact testing (Benjamini-Hochberg FDR = 0.05).

### Immune fluorescence and analysis with IMARIS

Cells were fixed with 4% paraformaldehyde, permeabilized and blocked in 2% BSA/0.2% TritonX100/PBS for 30 min at RT and incubated with the primary antibody (Golgin-97, #13192, mTOR #2983, RAB5 #3547, RAB7 #9367, Cell Signaling; LAMP2, sc-18822, Santa Cruz) in 1% BSA/0.1% TritonX100/PBS for 1 h at RT. After subsequent incubation with Alexa Fluor 488-conjugated secondary antibodies in 0.05% TritonX100/PBS for 45 min at RT, cells were mounted using Fluoroshield (ImmunoBioScience) containing 1 ng/µl DAPI. For Golgi visualization and mTOR-LAMP2 co-localization z-stack images (0.1 µm intervals) were acquired with 63x magnification using an inverted, confocal laser scanning microscope (LSM880+ Airyscan, Zeiss) and the ZEN software keeping imaging parameters constant for all slides. 3D image analysis of subcellular structures and cell size was performed with Imaris 9 (Bitplane), using the in-built cell and surface detection algorithm for cell size and Golgi structures, respectively. Golgi structures were detected utilizing background subtraction, structures below 400 volume pixel were excluded from the analysis. Endosomal vesicles were analyzed as RAB5- and RAB7-positive dots using the Imaris 9 spot detection algorithm. Default settings were applied, only the estimated size was set to 0.5 µm in diameter. The detection intensity threshold was set to automatic thresholding. LAMP2 and mTOR co-localization was analyzed using the Imaris 9 co-loc algorithm, setting intensity threshold for the analysis manually to exclude background fluorescence.

### Mass spectrometry-based proteomics of neonatal plasma

All chemicals from Sigma unless otherwise noted (Sigma-Aldrich Chemie GmbH, Munich, Germany). Plasma samples were centrifuged for 1 min at 2,000xg. Of the supernatant, 50 µg of protein per plasma sample were subjected to in-solution digestion with the iST 96x sample preparation kit (Preomics GmbH, Martinsried, Germany) according to manufacturer’s recommendations (3 h digestion).

Peptides were separated with a Dionex Ultimate 3000 RSLC nano HPLC system (Dionex GmbH, Idstein, Germany). 10 µg peptides were dissolved in 10 µL 0.1% formic acid (FA, solvent A) and 1 µl was injected onto an analytical column (400 mm length, 75 µm inner diameter, ReproSil-Pur 120 C18-AQ, 3 µm). Peptides were separated during a linear gradient from 5% to 35% solvent B (90% acetonitrile, 0.1% FA) at 300 nl/min over a 180 min. The LC was coupled to an Orbitrap Fusion Lumos mass spectrometer (Thermo Fisher Scientific, Bremen, Germany). Data-independent acquisition was performed with the following scan parameters: 47 windows of 15 Da plus 0.5 Da overlap covering m/z 399.5 to 1105.5. Isolated ions were fragmented with higher energy collision induced dissociation (HCD) with 22%, 27%, and 32% stepped collision energy. Fragments were detected in the Orbitrap detector (profile mode) with a resolution of 30,000 in the range of 200-1800 m/z. AGC target was 500,000, maximum injection time 50 ms. Every 3 s an MS1 scan was recorded in the range of 350-1500 m/z, resolution = 120,000.

Data processing was performed with DIA-NN 1.8.1 (Demichev et al. 2020) in library-free mode based on the Uniprot mouse reference proteome with isoforms (2023_03). The following parameters were applied: tryptic cleavage with one missed cleavage, variable modification of methionine by oxidation, acetylation of protein N-terminus, static modification of cysteine by carbamidomethylation, output filtered at 1% FDR.

The statistical analyses of the DIA-NN precursor ion quantities were carried out in the R environment (R version 4.2.3) (R Foundation for Statistical Computing 2018) using an in-house developed workflow. Quantities with more than 65% missing values were removed. The data were variance-stabilized and transformed using the VSN package version 3.64.0 (Huber et al. 2002) and missing values were imputed using the method “v2-mnar” of the msImpute package version 1.6.0 (Hediyeh-Zadeh et al. 2023). The data were aggregated on protein level using Tukey’s median polish method. The statistical analysis to identify differentially abundant proteins was performed with the limma package version 3.52.4 (Ritchie et al. 2015). To account for correlations, present in the same litter of mice, litter was modelled as random effect in the statistical analysis while using litter as blocking parameter in the function duplicateCorrelation. The resulting p-values of the statistical contrast between the knockout and wildtype condition were adjusted for multiple testing and the false discovery rates (FDR) were calculated by the Benjamini-Hochberg method.

### Statistics

Student’s *t* test (n ≥ 6 and Gaussian distribution), Mann-Whitney test (n ≤ 5 or no Gaussian distribution) or one-way ANOVA were calculated with GraphPad Prism software (San Diego, CA, USA). Values of p < 0.05 were considered significant. Results are given as mean±SD or in a scatter plot with individual data points.

## Acknowledgements

We thank members of the Kolanus and Meissner labs for technical help and useful discussions, and we thank Jörg Höhfeld for comments on the manuscript.

We would like to thank the Analytical Proteomics Core Facility and the *Core Unit* for *Bioinformatics* Data *Analysis* of the Medical Faculty at the University of Bonn, namely Marc Sylvester, Andreas Buness, and Farhad Shakeri, for providing support and instrumentation funded by the Deutsche Forschungsgemeinschaft (DFG, German Research Foundation) – including Projektnummer 386936527.

The work was funded by the Deutsche Forschungsgemeinschaft (DFG, German Research Foundation) under Germany’s Excellence Strategy-EXC2151-390873048 to W Kolanus and DFG FOR 2743: Mechanical Stress Protection-388932620. Further, we would like to thank the TRR237 for providing support and resources for the development of the spatial proteomics analysis, funded by the Deutsche Forschungsgemeinschaft (DFG, German Research Foundation) – Projektnummer 369799452).

This article is subject to MI’s Open Access to Publications policy. MI lab heads have previously granted a nonexclusive CC BY 4.0 license to the public and a sublicensable license to HHMI in their research articles. Pursuant to those licenses, the author-accepted manuscript of this article can be made freely available under a CC BY 4.0 license immediately upon publication.

## Author Contributions

Conceptualization: C.K., B.J., W.K.

Methodology: C.K., B.J., F.S., S.K.

Investigation: C.K., B.J., F.S., S.K.

Writing - original draft: C.K., B.J.

Writing - review and editing: C.K., B.J., W.K., S.B., F.S., F.M.

Supervision: W.K., C.K., B.J.

Funding acquisition: B.J., W.K.

## Competing Interests Statement

The authors declare no competing interests.

## Data Availability

Proteomics data will be available on ProteomeXchange and the partner platform PRIDE soon.

## Supplementary Materials

Supplementary Materials and Methods

Figs. S1 to S6

Supplementary References

## Notes

### Competing Interest Statement

The authors have declared no competing interest.

