## Supplementary Information for "Cytohesin-2 is essential for the survival of mice and regulates Golgi volume and function"

### Supplementary Methods

#### Tissue-specific *Cyth2*-deficient mice

Tissue-specific *Cyth2*$^{-/-}$ mice were generated by breeding *Cyth2*$^{+/-}$ mice with Flp-deleter mice [B6.SJL-Tg(ACTFLPe)9205Dym/J, (Rodríguez et al. 2000)] to remove the transcriptional stop between two FRT-sites behind exon five. The following Cyth2 conditional mice were bred with nestin-Cre mice [brain-specific, (Tronche et al. 1999)], myogenin-Cre mice [skeleton muscle-specific (Li et al. 2005)], alpha-myosin heavy chain mice [heart muscle-specific, (Agah et al. 1997)] and albumin-Cre [liver-specific, (Kellendonk et al. 2000)] mice, respectively Genotyping of adult mice was done from tissue of ear labeling.

#### H&E-staining of heart tissue

PFA-fixed paraffin-embedded murine heart cross-sections were deparaffinized by in-cubation in xylol twice for 5 min. Next, the sections were rehydrated by a descending ethanol dilution series (100% (2x, 3 min), 96%, 70%, 50% (2 min each), and dipped in distilled water. Cross-sections were stained in haematoxylin for 1 min and washed in cold running tap water for 10 min. Afterwards, the sections were counterstained with 0.5% eosin for 2 min, excess dye was rinsed in cold running tap water. Cross-sections were dehydrated in an ascending ethanol dilution series [50%, 70%, 96%, 100% (2x)] and cleared in xylol twice for 2 min. Sections were then mounted with Entellan™ (Merck), dried under the fume hood and imaged by bright-field microscopy using a Olympus SZX10 (Olympus).

#### LysoTracker staining of C2 myoblasts

WT and Cyth2-deficient C2 myoblasts were seeded onto Ibidi-Slides (35 mm IbidiTreat) and incubated overnight under standard culture conditions. Afterwards, the cells were treated with 250 nM LysoTracker™ Green DND-26 (ThermoScientific) for 45 min in fresh medium, followed by three washing steps in live-cell imaging solution (HBSS, 5% FCS, 20 mM HEPES pH7.4) and imaging with a laser scanning confocal microscope. At least 20 cells per condition were imaged, analysis was performed using the ImageJ Analyze particles module identifying spots of 0.2-1.5 µm diameter.

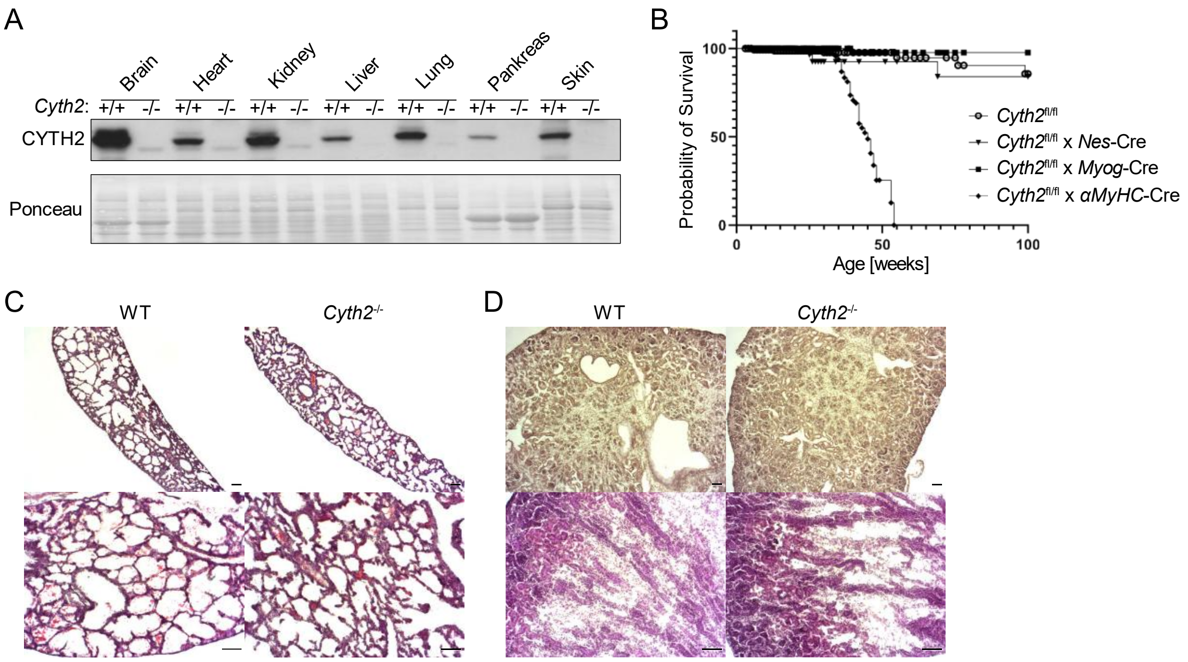

**Supplementary Figure 1: Tissue-specific analysis of *Cyth2* cannot explain the full-knockout lethality of *Cyth2*$^{-/-}$mice**

Conditional knockout of *Cyth2* in a variety of tissues could not pheno-copy full-knockout lethality. **A)** Exemplary Western Blot analysis of CYTH2 expression in neonatal organs (as indicated) of WT and knockout mice. Ponceau staining served as a loading control. **B)** Kaplan-Meier graph depicting the survival probability of organ-specific *Cyth2* knockout mice for brain, skeletal muscle, and heart compared to floxed *Cyth2* allele controls (as indicated by the respective Cre-breedings). Each symbol represents an individual mouse (n≥73). **C-D)** Hematoxylin-Eosin staining of **C)** kidney and **D)** lung cryosection from WT and *Cyth2* knockout mice six hours after birth. Scale bars indicate 50 µm.

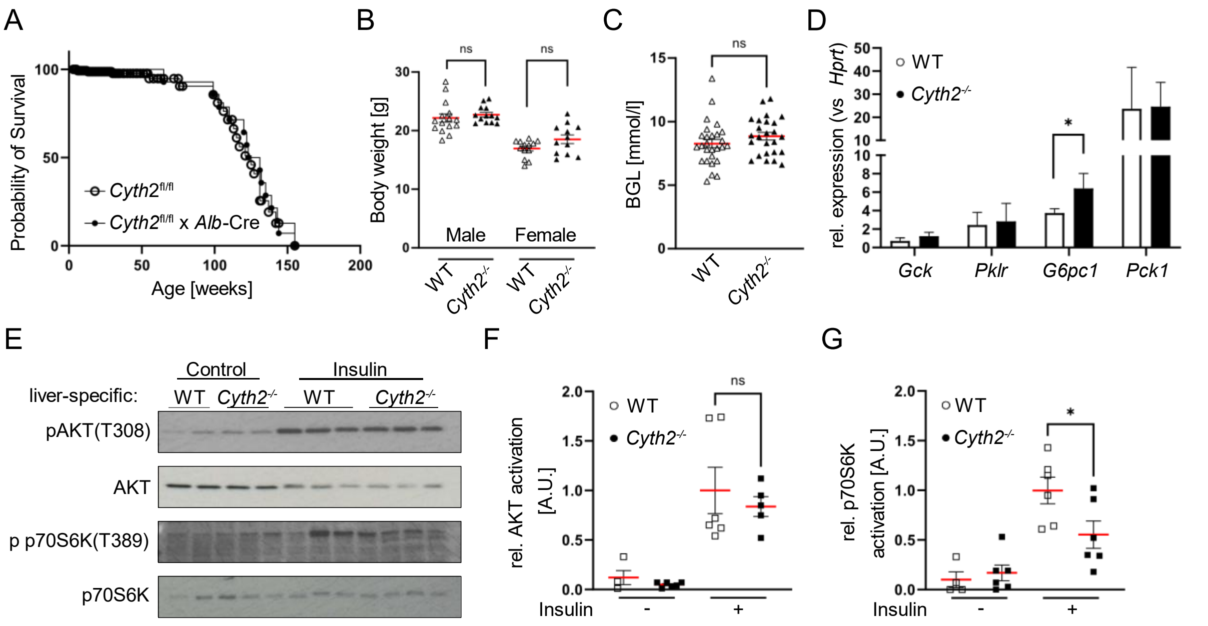

**Supplementary Figure 2: Liver-specific loss of *Cyth2* points towards a metabolic function of the small ARF-GEF.** Liver-specific knockout of *Cyth2* does not phenocopy full-knockout mice and indicates a metabolic impact of *Cyth2*. **A)** Kaplan-Meier graph depicting the survival probability of liver-specific *Cyth2* knockout mice compared to floxed controls. Each symbol represents an individual mouse (n≥14). **B)** Body weight of adult liver-specific *Cyth2* knockout mice compared to respective WT controls (n≥12).**C)** Blood glucose levels of adult WT and liver-specific *Cyth2* knockout mice (n≥25). **D)** Relative mRNA expression of glucose homeostasis genes in hepatic tissue of WT and liver-specific *Cyth2*-deficient adult mice, normalized to *Hprt* (n=4). **E-G)** Insulin signaling in hepatic tissue from adult liver-specific *Cyth2* knockout mice analyzed by Western Blot of phosphorylated (activated) AKT and S6K. **E)** Representative Western Blots of phosphorylated AKT and S6K in hepatic tissue from mice treated with or without insulin. Total AKT and S6K served as loading controls. Organs were isolated 10 min after insulin injection (i.p.; 0,75 U/kg human insulin) and snap-frozen in liquid nitrogen. **F-G)** Semi-quantitative analysis of AKT and S6K phosphorylation. Phosphorylated signals were normalized to total protein signals. Data are represented as mean±SD; each symbol represents an individual animal. Significance was tested using unpaired t-test or Mann-Whitney test (*$p < 0.05$, ns = not significant).

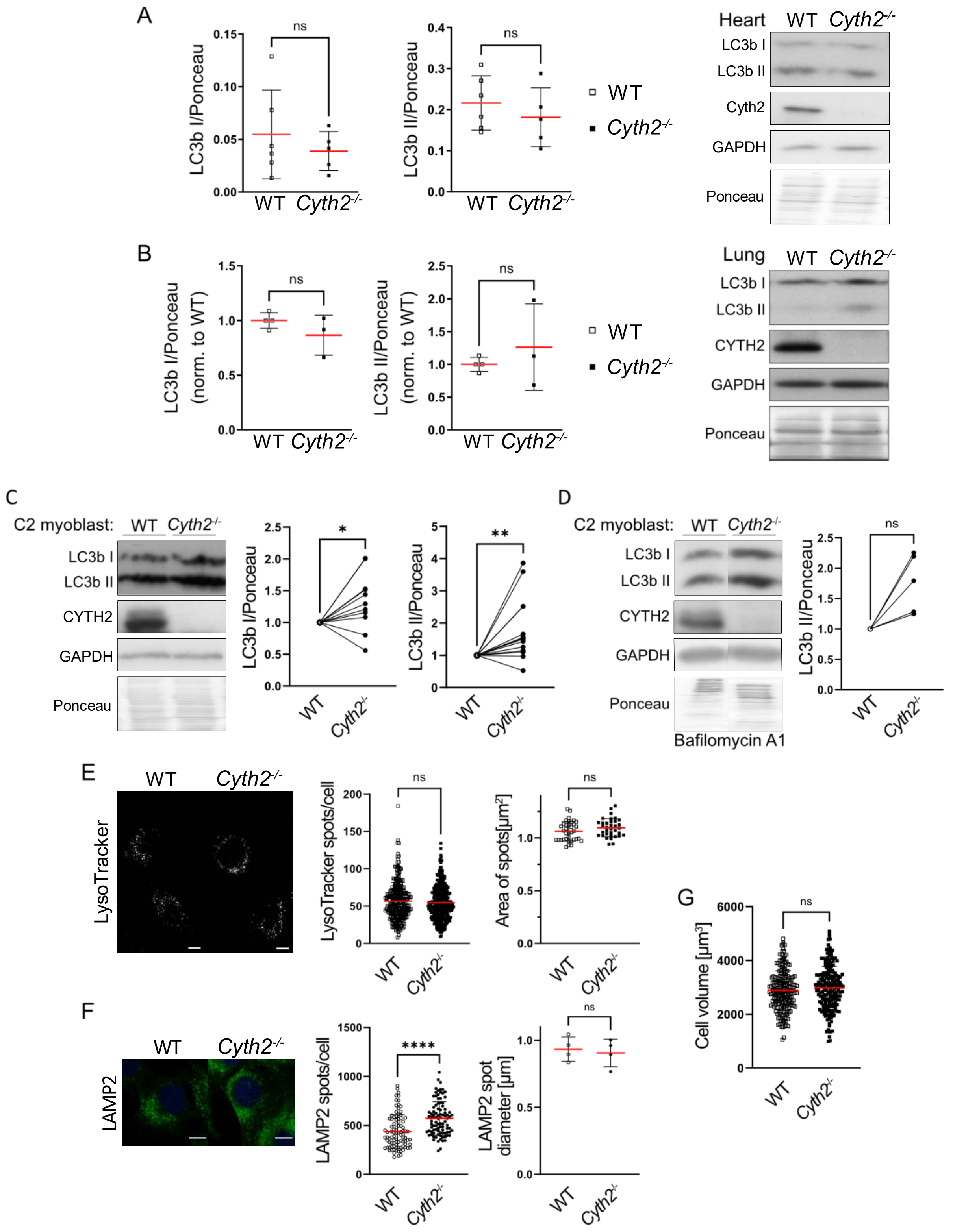

**Supplementary Figure 3: *Cyth2* deficiency does not dramatically affect autophagy in neonatal tissue or C2 myoblasts**

*Cyth2* knockout tissues and cells have unaltered or even slightly increased autophagic activity. **A)** Western Blot analysis of LC3b I and II expression levels in neonatal heart tissue of WT and knockout mice. Representative blots are shown on the right. Ponceau staining served as a loading control (n≥5). **B)** Western Blot analysis of LC3b I and II expression levels in neonatal lung tissue of WT and knockout mice. Representative blots are shown on the right. Ponceau staining served as a loading control (n≥3). **C)** Western Blot analysis of steady state LC3b I and II expression levels in WT and *Cyth2*-deficient C2 myoblasts, including representative blots. Cells were untreated and harvested after two days of standard culture conditions. Ponceau staining served as a loading control, data were normalized to the WT control (n≥11). **D)** Western Blot analysis of LC3b I and II expression levels in WT and *Cyth2*-deficient C2 myoblasts with blocked lysosomal acidification, including representative blots. Cells were treated for 2-4 h with Bafilomycin A1 after two days standard culture conditions. Ponceau staining served as a loading control, data were normalized to the WT condition (n=5). **E)** Fluorescence analysis of LysoTracker-stained WT and *Cyth2*-deficient C2 myoblasts after two days standard culture conditions (steady state). LysoTracker-positive vesicles were counted and their size measured. **F)** Fluorescence analysis of LAMP2-stained WT and *Cyth2*-deficient C2 myoblasts after two days standard culture conditions (steady state). LAMP2-positive vesicles were counted and their diameter measured **G)** Myoblasts were surface-stained to measure the volume of WT and *Cyth2* knockout cells, serving as a control for increased organelle counts. Representative cells are depicted, data are represented as mean±SD; each symbol represents an individual cell or the average of one image acquired (n≥274; n≥35). Significance was tested using unpaired t-test, Wilcoxon or Mann-Whitney test (*$p < 0.05$, **$p < 0.01$, ns = not significant). Scale bars represent 10 µm.

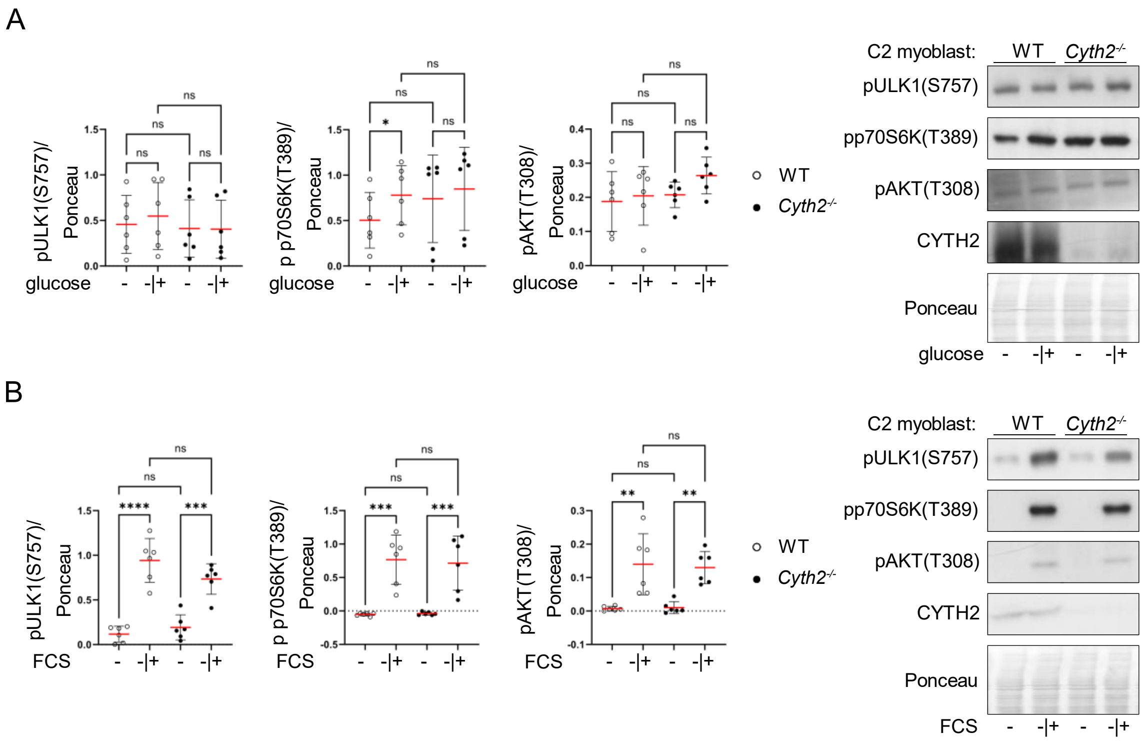

**Supplementary Figure 4: *Cyth2*-deficiency does not affect the response of C2 myoblasts towards glucose or FCS.**

**A-B)** Western Blot analysis of phosphorylated ULK1, p70S6K and AKT in C2 myoblasts after deprivation (-) of **A)** glucose (and sodium pyruvate) or **B)** FCS for two hours and subsequent restimulation (-|+) for two hours. Left: Average signal intensity (red line±SD), normalized to total protein loaded as determined by Ponceau staining. Data were pooled from three independent experiments including two pairs of WT and *Cyth2* knockout clones. Each dot represents one measurement (n=6). Significance was tested using 2way-ANOVA (*p < 0.05; **p < 0.01; ***p < 0.001; ****p < 0.0001; ns = not significant). Right: representative immunoblot analyses.

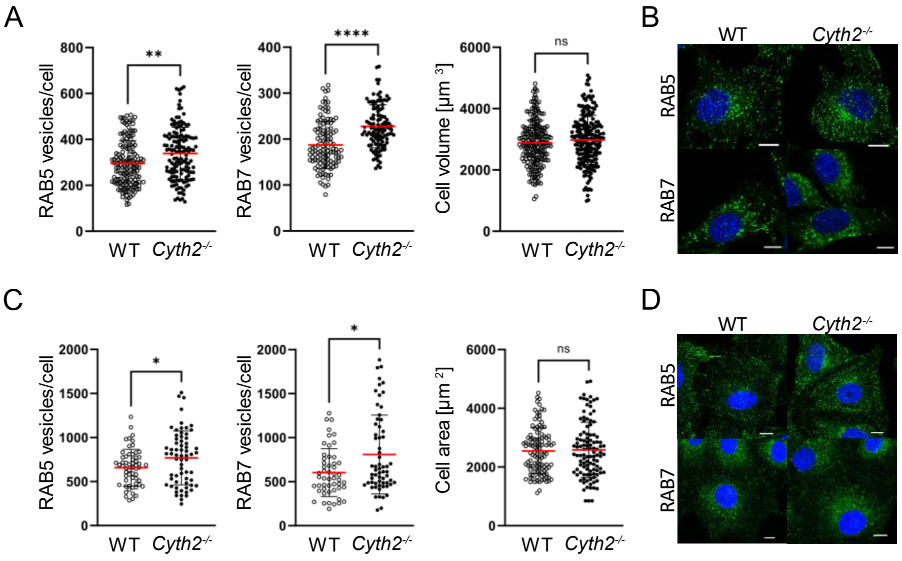

**Supplementary Figure 5: *Cyth2* deficiency increases the numbers of endosomal compartments in C2 myoblasts.**

**A-B)** Immunofluorescence analysis of RAB5 and RAB7 in WT and *Cyth2*-deficient C2 myoblasts. The counts of **A)** RAB5-positive vesicles/cells and RAB7-positive vesicles per cell were analyzed, cells size was measured as a control by surface staining. **B)** Representative images of stained cells. Data were pooled from independent experiments (n≥6) and depicted as mean±SD, each dot representing an individual cells (n≥51). Significance was tested using Mann-Whitney test (*p < 0.05; **p < 0.01; ***p < 0.001; ****p < 0.0001; ns = not significant). Scale bars represent 10 µm.

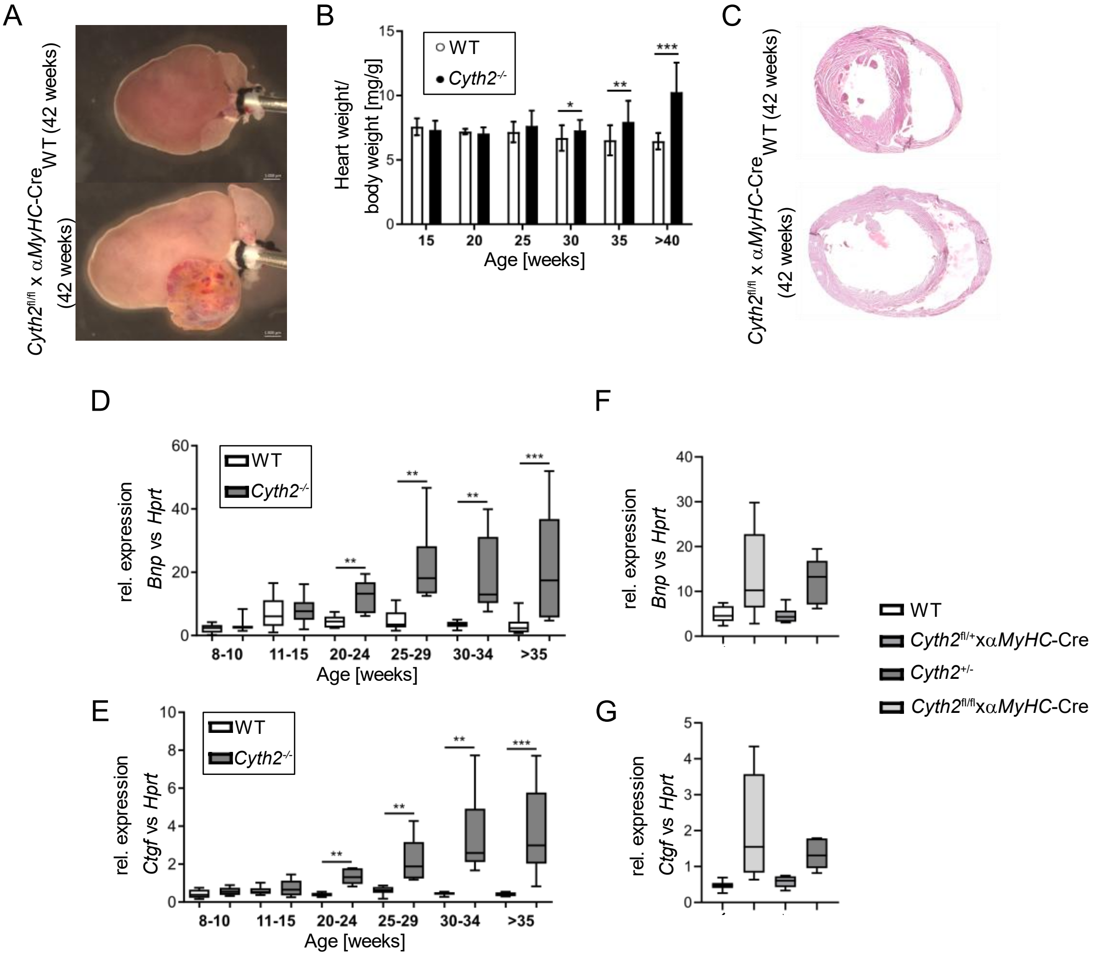

**Supplementary Figure 6: Description of the heart-specific *Cyth2*$^{-/-}$ mousline developing cardiac hypertrophy.**

**A)** Representative images of 42-week-old hearts isolated form *Cyth2*$^{fl/fl}$ and *Cyth2*$^{fl/fl}$ x *αMyHC*-Cre mice, fixed on a canula. **B)** Heart-to-body weight ratio of *Cyth2*$^{fl/fl}$ and *Cyth2*$^{fl/fl}$ x *αMyHC*-Cre mice over time (n≥5). **C)** Representative H&E-stained cross-sections of 42-week-old hearts of *Cyth2*$^{fl/fl}$ and *Cyth2*$^{fl/fl}$ x *αMyHC*-Cre mice. **D-E)** Relative mRNA expression of **D)** *Bnp* and **E)** *Ctgf* in ventricle tissue of *Cyth2*$^{fl/fl}$ and *Cyth2*$^{fl/fl}$ x *αMyHC*-Cre mice over time, *Hprt* serving as the housekeeping gene (n≥6). **F-G)** Relative *Bnp* and *Ctgf* mRNA expression as in E-F) in ventricle tissue isolated from aged (20-33 weeks) WT, *Cyth2*$^{fl/+}$ x *αMyHC*-Cre, *Cyth2*$^{+/-}$ and *Cyth2*$^{fl/fl}$ x *αMyHC*-Cre mice (n≥7). Significance was tested using students' t-test or Mann-Whitney. (*p < 0.05; **p < 0.01; ***p < 0.001; ns = not significant).

**Supplementary Table 1. Primers for Genotyping and qPCR**

| Target gene | Primer name | Sequence (5’-3’) |
|---|---|---|
| Genotyping | | |
| *Cyth2* (murine) | Pscd2 WT1 screen for | CAGAAATGCCAGGGCTTTCTCAGC |
| | Pscd2 WT1 screen rev | GCATAGGTTTCAGGGCTGGAAAACAC |
| | Pscd2 FRT screen rev | CGGAAGGAATGCCCAGCCAAAAT |
| Cre recombinase | CRE for | CCGGTCGATGGAGTGA |
| | CRE rev | GGCCCAAATGTGGATA |
| qPCR | | |
| mouse *G6pc1* | mG6pc1 for | AGC TGA ACG TCT GTC TGT CC |
| | mG6pc1 rev | TTC TCC AAA GTC CAC AGG AG |
| mouse *Gck* | mGck for | GAA CAA CAT CGT GGG ACT TC |
| | mGck rev | AGC TCC ACA TTC TGC ATC TC |
| mouse *Pklr* | mPklr for | CAT TGC TGT GAC TCG TTC TG |
| | mPklr rev | CAC AAT CAC CAG ATC ACC AA |
| mouse *Pck1* | mPck1 for | TGC CGG AAG AGG ACT TTG AG |
| | mPck1 rev | CAC TTG ATG AAC TCC CCA TC |

**Supplementary Table 2. sgRNAs for CRISPR knockout of *Cyth2***

| Target gene | Sequence (5´-3´) | sgRNA library |
|---|---|---|
| Mouse *Cyth2* | TTTGCAGTAAGACCTTGCAG | Brie |
| Human *CYTH2* | GCTCAGTGAAGCCATGAGCG | Brunello |

**Supplementary Table 3: CYTH2-overexpression constructs for rescue experiments**

All inserts were verified by sequencing (GATC Eurofins…).

| Vector backbone | Gene | Mutation | 5´Tag |
|---|---|---|---|
| pN1 | *Cyth2*-2G | None | RFP |
| pN1 | *Cyth2*-2G | E156K | RFP |
| pN1 | *Cyth2*-2G | DCC (2-46AS) | RFP |
| pN1 | *Cyth2*-3G | None | RFP |
| pN1 | *Cyth2*-3G | E156K | RFP |
| pN1 | *Cyth2*-3G | DCC (2-46AS) | RFP |
